# Single Cell Transcriptomics-Informed Induced Pluripotent Stem Cells Differentiation to Tenogenic Lineage

**DOI:** 10.1101/2023.04.10.536240

**Authors:** Angela Papalamprou, Victoria Yu, Wensen Jiang, Julia Sheyn, Tina Stefanovic, Angel Chen, Chloe Castaneda, Melissa Chavez, Dmitriy Sheyn

**Author notes:** Correspondence: Dmitriy Sheyn, PhD Assistant Professor Board of Governors Regenerative Medicine Institute Department of Orthopedics Department of Surgery Department of Biomedical Sciences Cedars-Sinai Medical Center 8700 Beverly Blvd. AHSP A8308 Los Angeles, CA, 90048, USA.

## Abstract

During vertebrate embryogenesis, axial tendons develop from the paraxial mesoderm and differentiate through specific developmental stages to reach the syndetome stage. While the main roles of signaling pathways in the earlier stages of the differentiation have been well established, pathway nuances in syndetome specification from the sclerotome stage have yet to be explored. Here, we show stepwise differentiation of human iPSCs to the syndetome stage using chemically defined media and small molecules that were modified based on single cell RNA-sequencing and pathway analysis. We identified a significant population of branching off-target cells differentiating towards a neural phenotype overexpressing Wnt. Further transcriptomics post-addition of a WNT inhibitor at the somite stage and onwards revealed not only total removal of the neural off-target cells, but also increased syndetome induction efficiency. Fine-tuning tendon differentiation *in vitro* is essential to address the current challenges in developing a successful cell-based tendon therapy.

## Introduction

Tendons are necessary to support body movement by transferring forces between muscle and bone. Unfortunately, tendon injuries are quite prevalent in athletes and in the aging population, comprising 45% of musculoskeletal consultations in the US alone.^1^ Tendons are structurally complex tissues, and their low vascularity and cellularity are contributing factors to their poor regenerative capacity.^2^ The current standard of treatments for severe tendon injury includes conservative approaches and/or surgical intervention.^1^ However, the resulting prolonged rehabilitation, muscle weakness, and high re-injury rates dramatically limit patient outcomes.^3^ Development of a stem cell-mediated solution could improve patient outcomes, yet current approaches fall short in terms of tissue biomechanical performance and tissue organization.^4–6^

Cell-based therapies have shown potential for tendon repair.^7^ Autologous tendon progenitors and Mesenchymal Stromal Cells (MSCs) have been explored as potential cell sources, however, their limitations for clinical application are associated with their heterogeneity and the need for *in vitro* expansion for obtaining clinically relevant cell numbers. Cell expansion *in vitro* can result in phenotypic drift and subsequent functional loss, in addition to low proliferative ability.^8^ Utilizing tenocytes differentiated from induced pluripotent stem cells (iPSCs) holds promise due to their unparalleled developmental plasticity, unlimited self-renewal capacity, and the potential scalability for an off-the-shelf cell source application. Well-established protocols have been developed for the differentiation from pluripotent stem cells to different musculoskeletal cell types including chondrocytes and osteoblasts.^9–12^ Recent studies have successfully differentiated tenocytes using mouse iPSCs and Embryonic Stem Cells.^13–16^ Though, recent work has shown a divergence in developmental processes between mouse and human embryos and subsequent differences in developmental cues on tenogenic differentiation between species.^17–20^ This highlights the importance of researching the developmental cues of human cells. Furthermore, other studies have either not reported induction efficiency or showed limited syndetome-like/tenocyte induction.^13,21^ Deriving a homogenous population of tenocytes from iPSCs with high efficiency continues to be a challenge, largely in part to the limited understanding of their developmental origins and differentiation path from multipotential precursors.

In vertebrate embryogenesis, tendons arise from mesodermal cells of different origin. Axial, limb, and cranial tendons originate from paraxial mesoderm, lateral plate mesoderm or neural crest (NC), respectively. Despite differential cell and tissue interactions between the three regions, the major molecular regulators of tendon differentiation are shared between all three.^22,23^ BMP, TGFβ, Activin/Nodal, FGF and WNT signaling pathways regulate mesoderm induction from pluripotent cells. Along the mediolateral axis of the embryo the balance between BMP and WNT signaling gradients specify mesodermal subtypes.^24,25^ The paraxial or presomitic mesoderm (PSM) is derived from Neuromesodermal Progenitors (NMPs) that are bipotential and able to differentiate into ectodermal and mesodermal lineages.^24^ Recent *in vitro* studies demonstrated that during mesoderm specification, WNT controls the allocation to PSM and represses lateral plate mesoderm.^25,26^ Somites (SM) are derived from the anterior PSM following dynamic morphogenetic cyclic signaling, involving Notch, WNT and FGF.^24^ Somite progenitors are multipotential and are further specified by signaling molecules from the surrounding tissues. Sonic hedgehog (SHH) is secreted from the notochord and the neural tube and it has been demonstrated as a crucial signaling modulator in sclerotome (SCL) specification of somites combined with low levels of WNT and BMP.^24,25^ Murine and avian embryonic development studies concluded that combined SHH activation and BMP inhibition may be sufficient for SCL induction.^24^ However, it was later shown *in vitro* that WNT directly antagonizes SHH and induces somites to dermomyotome.^25^ Since SCL is in contact with different cell populations and thus diverging signals, it gives rise to different cell populations along the three major patterning axes. SCL develops dorsally into the syndetome (SYN), the precursor of tendons of the body trunk, which is marked by the expression of the transcription factor scleraxis (SCX).^24^ Recent studies have indicated that FGF, BMP and TGFβ are orchestrating syndetome differentiation from sclerotome.^15,16,27^ A few groups have differentiated mouse iPSCs^15^ and human iPSCs^27^ to SYN through SCL and have pinpointed the primary role of BMP and TGFβ signaling. However, the nuances of WNT signaling in SYN specification have yet to be explored.

In this study, we sought to fine-tune induction to SYN by further elucidating the signaling pathways involved with tenocyte formation using single cell transcriptomics. SYN was induced from GMP-ready human iPSCs by stepwise differentiation through PSM-derived SM cells using chemically defined media. At every stage of the induction, we monitored known markers for each developmental step and conducted further characterization with single-cell RNA sequencing (scRNA-seq) and immunostaining. ScRNA-seq analysis revealed off-target differentiation towards a neural phenotype, with WNT family members being identified as crucial to the generation of neural by-products. We hypothesized that WNT signaling inhibition would block cell fate bifurcation to “unwanted” states and drive induction towards a single path. Informed by single cell transcriptomics, we added the WNT inhibitor Wnt-C59 to the later stages of the differentiation, which improved final induction efficiency of the differentiated tendon progenitor population.

## Results

### Stepwise induction of iPSCs to syndetome-like cells using chemically defined media and small molecules *in vitro*

Human iPSCs were seeded at different densities and induced to PSM for 4 days with the GSK3 inhibitor (GSK3i) CHIR99021 leading to WNT pathway activation, combined with BMP and TGFβ inhibition and concurrent FGF activation (Fig. 1A). Gene expression analysis, immunofluorescent staining (IF) and flow cytometry for DLL1 levels were used to determine induction efficiency and establish the starting seeding density (Suppl. Fig. 1). A homogeneous population of DLL1+ cells was generated after 4 days when cells were seeded at ∼40-50 aggregates/cm^2^. Specifically, 96.2% DLL1^+^ cells were generated at day 4 (Suppl. Fig.1). In contrast, DLL1^+^ cells were shown to be 43.6% less (52.6% DLL1^+^ cells) when iPSCs were seeded at 2x concentration.

**Fig. 1.**
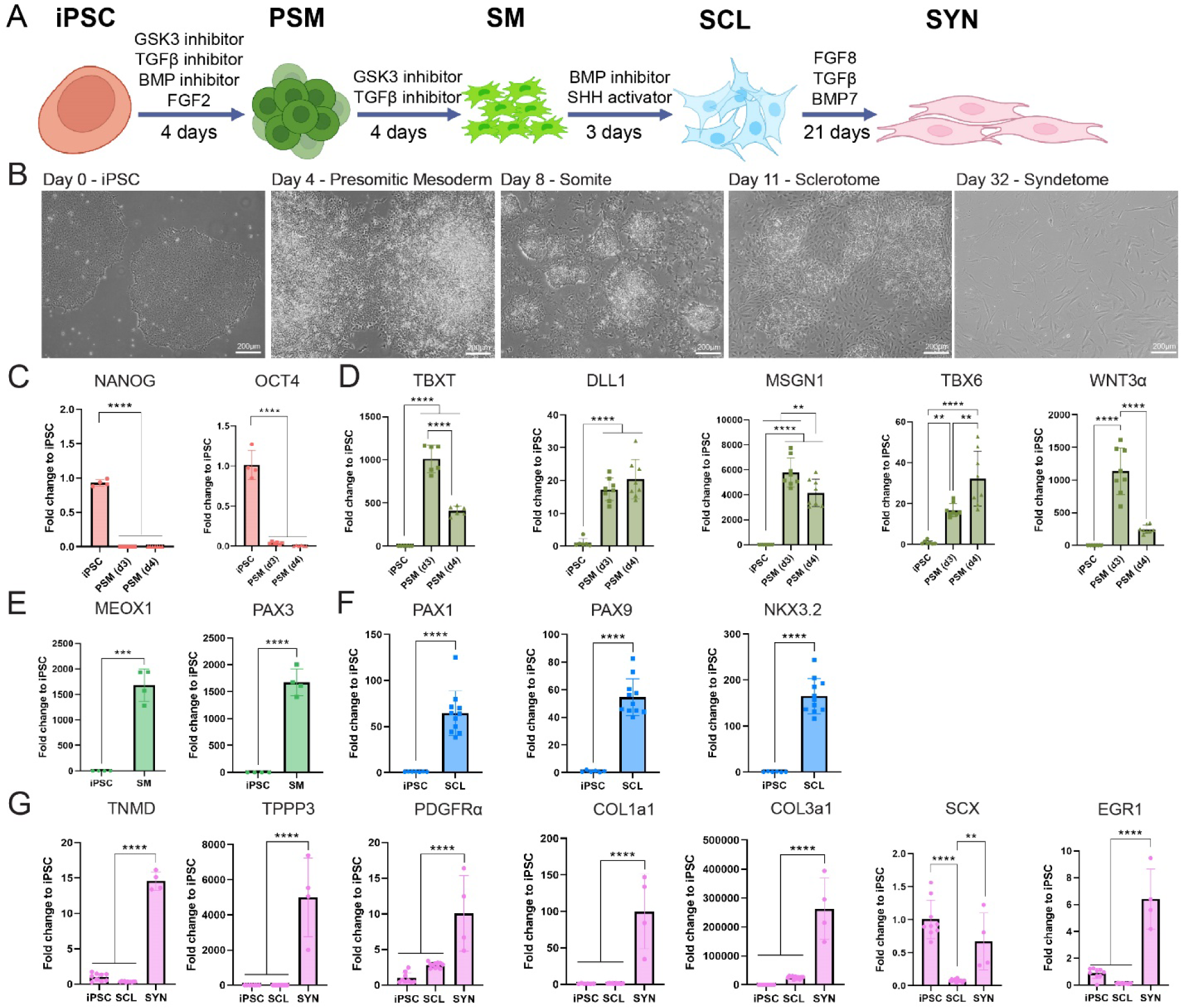
Stepwise induction of iPSCs to syndetome-like cells using chemically defined media and small molecules *in vitro.* **(A)** Schematic of iPSC to SYN stepwise induction using chemically defined media and small molecules. **(B)** Brightfield micrographs of cells going through the differentiation stages at. Scale bars represent 200μm. **(C)** Pluripotent markers were expressed in iPSCs and were downregulated in further stages. **(D-F)** Gene expression analyses for stage-specific markers with the 007i iPSC line: upregulation of early mesoderm markers at the PSM (n=8 replicates/group) **(D)**, somitogenesis at SM (n=8 replicates/group) **(E)**, sclerotome-related markers at SCL (n=12) **(F)**. **(G)** Tenogenic markers are significantly upregulated at the SYN stage (n=4) compared to iPSC (n=9) and SCL (n=12) stages. Differentiation experiments were repeated independently n=2 with the 007i line and n=2 with a second iPSC line that was later tested (83i); \**p<0.05, **p<0.01, ***p<0.001, ****p<0.0001*.

Cells exhibited distinct morphologies at different stages of chemical induction (Fig. 1B). On day 4, iPSC colonies displayed contraction, resulting in differentiating cells leaving the colonies at the periphery. By day 8, cells were completely dissociated from the colonies. On day 11, colonies looked contracted and regressed, and were surrounded by spindly cells. Lastly, by the end of the differentiation period, fibroblast-like cells had grown in size. However, during SYN induction we noticed a gradual attrition of cells, resulting in considerably lower cell numbers at the end of the process.

To further characterize cell differentiation at each stage, the expression of PSM, SM and SCL stage cell markers was examined with RT-qPCR (Fig. 1C-F) and IF (Fig. 2A-G). Additionally, markers associated with tendon progenitors and tenocytes were examined as an indication of SYN differentiation (Fig. 1G, Fig. 2H-I). Pluripotency markers (OCT4, NANOG) were significantly downregulated at PSM compared to iPSC and were not detected at subsequent stages (Fig. 1C). PSM marker genes DLL1, TBX6, WNT3A and MSGN1^24,25^ were significantly upregulated at day 3 and day 4 of GSK3i treatment and downregulated after 4 more days of TGFβ inhibition and WNT activation of DLL1^+^ cells (Fig. 1D). Mesodermal T-box transcription factors TBX1 and TBX6^28^ were detected at PSM stage (Fig. 2C) but not at SM stage (Fig. 2C-D). SM markers MEOX1, and PAX3^24,25^ were upregulated at day 8, defining the somitogenesis stage (Fig. 1E, Fig. 2C-E). Following 3 days of BMP inhibition and SHH pathway activation of SM stage cells, SCL markers (NKX3.2, PAX1 and PAX9)^24,25^ were significantly upregulated at day 11 (Fig. 1F). PAX1 and PAX9 were observed in the entire cell population at SCL (Fig. 2G). PDGFRA and TPPP3^29^ markers were significantly upregulated at the SYN stage (Fig. 1G). Interestingly, SCX, a prominent tenogenic transcription factor,^30^ was significantly downregulated at the SCL stage compared to iPSC, but upregulated during the differentiation from SCL to SYN. Further, COL1A1, COL3A1 and TNMD^31^ were significantly upregulated at the last stage of SYN induction (Fig. 1G). IF staining confirmed the presence of SCX^+^COL1^+^ and TNMD^+^COL1^+^MKX^+^ cells at the end of the induction (Fig. 2I). However, expression of tenogenic markers was heterogeneous in different cells (Fig. 2I). Additionally, we noticed gradual attrition of cells after SCL stage, resulting in considerably lower cell numbers by the end of the differentiation.

**Fig. 2.**
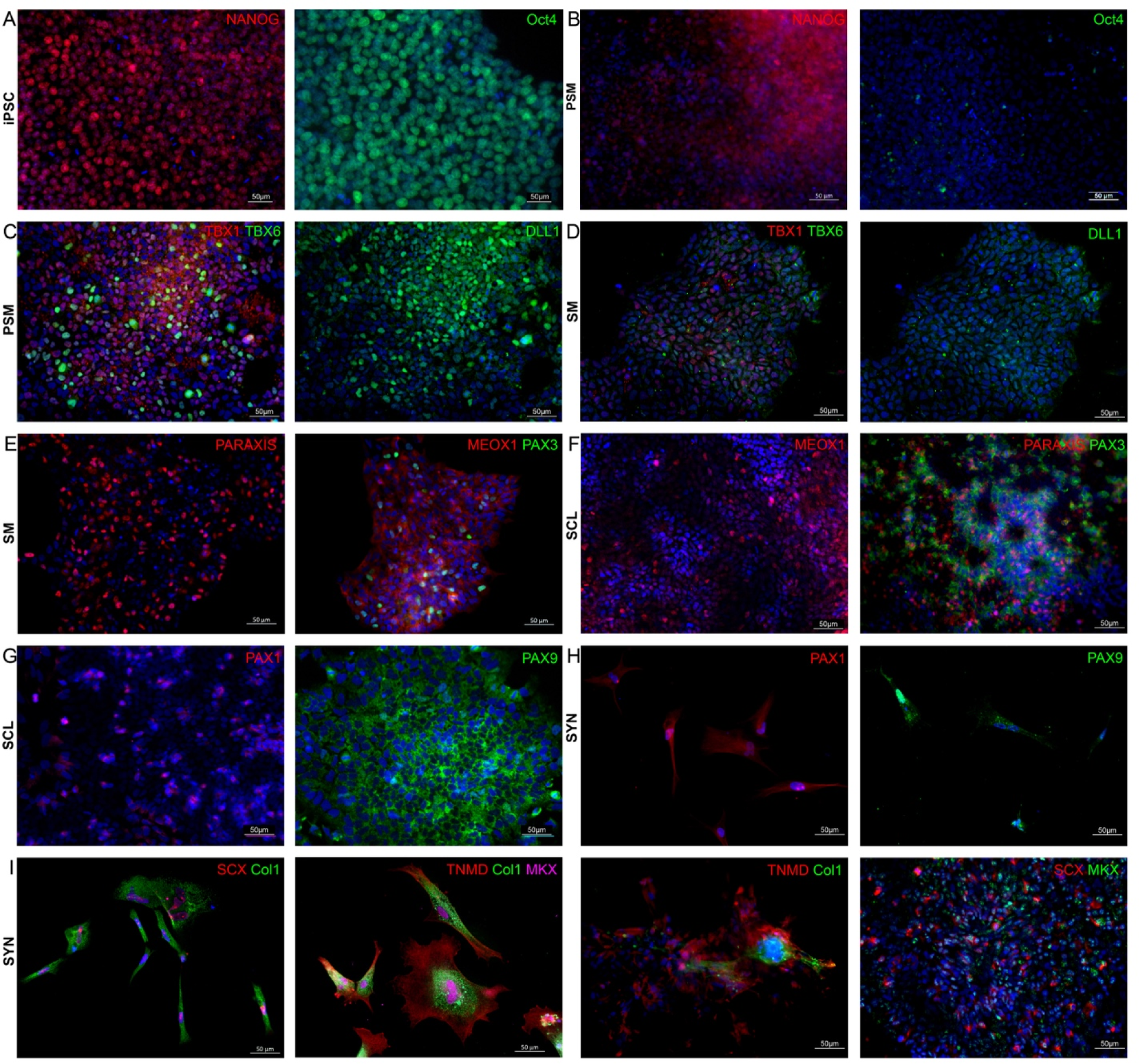
Immunofluorescence co-staining for stage-specific markers for confirmation of protein expression at each induction stage. **(A-B)** OCT4 and NANOG were expressed in iPSCs but were downregulated at PSM. **(C-D)** Early mesoderm markers TBX1, TBX6 and DLL1 at PSM vs. SM. **(E-F)** Somitogenesis markers PARAXIS, MEOX1, and PAX3 at SM vs. SCL. **(G-H)** sclerotome-related markers PAX1 and PAX9 at SCL vs. SYN. **(I)** Tenogenic markers SCX, COL1, TNMD, MKX co-expression at the end of induction to SYN. Nuclei were stained with DAPI (blue). Scale bars represent 50μm. Differentiation experiments were repeated independently n=2 with the 007i line and n=2 with a second iPSC line that was later tested (83i). IF staining was performed in n=3 technical triplicates.

### Single-cell RNA sequencing reveals cellular heterogeneity at the end of induction of iPSC to syndetome-like cells

Even though gene expression and IF showed expression of early tendon markers by the end of differentiation using two iPSC lines (Suppl. Fig. 4A), cell heterogeneity was observed with IF (Fig. 2). Further, SYN differentiation efficiency was suboptimal, resulting in considerable cell attrition by the end of the induction process (Fig. 1B, Fig. 2G-I). To further explore the observed cellular heterogeneity, we sought to investigate the cell transcriptome at each stage (iPSC, PSM, SM, SCL, and SYN) at a single cell level using the GMP-ready line 007i and scRNA-seq analysis. The dimensional reduction based on the unsupervised UMAP method sorted cells into 11 clusters that could be classified as 6 cell subpopulations (Fig. 3A). Raw counts of cell numbers for each cluster and stage are shown in Suppl. Table 1 and normalized proportions per cluster in Suppl. Fig. 2. Feature plots and dot plots showed upregulation of stage-specific genes (Fig. 3B-C). That is, pluripotency markers (OCT4, NANOG, SOX2)^32^ were expressed at iPSC and disappeared at the PSM stage, while PSM markers TBXT, TBX6, MSGN1, DLL1, DLL3^24,25,33^ were upregulated. SM markers (MEOX1, PAX3)^24,25^ and SCL markers (PAX1, PAX9, NKX3.2, SOX9)^24,25^ were upregulated in a stepwise manner. Tenogenic markers (SCX, MKX, TNMD, COL1A1, DCN, COL3A1)^31,34^ were found in SYN cluster C3 (Fig. 3B-D). These plots show that the SYN cluster accounted for 48.4% of the entire population (Suppl. Table 1 and Suppl. Fig. 2). Notably, two “side-arm” clusters, which became prominent in SCL, comprised of 18.3% of the cells collected at the SYN stage (Fig. 3A, Suppl. Table 1 and Suppl. Fig. 2). These were denoted as neuromesodermal progenitor – cranial cell populations hereafter referred to as NMP-C (DLL1^+^DLL3^+^NOTCH1^+^CRABP1^+^),^25,35,36^ and neural lineage, hereafter referred to as NL (NRN1^+^DCX^+^NNAT^+^),^37–39^ cell populations.

**Fig. 3.**
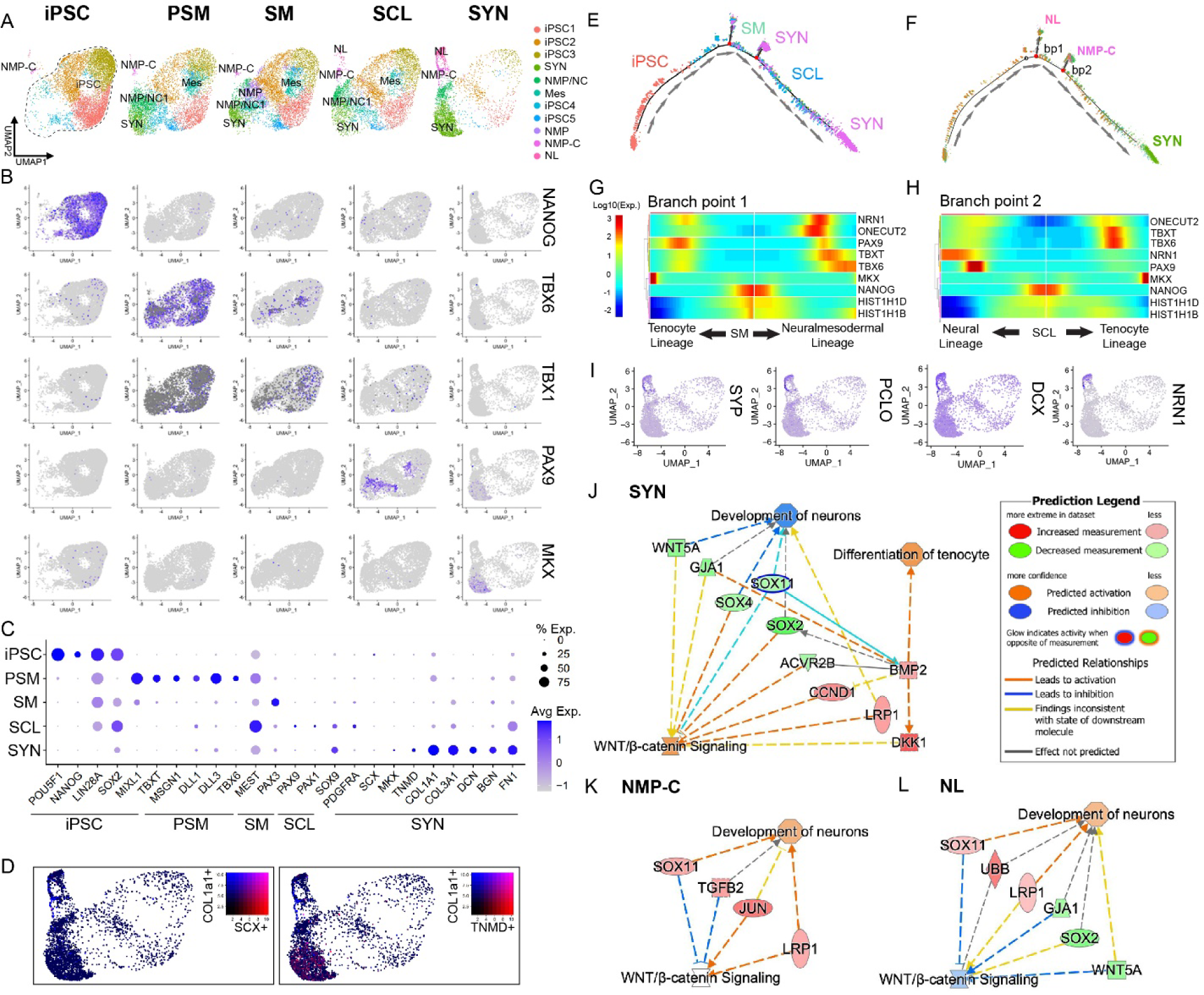
Single-cell RNA sequencing reveals cellular heterogeneity at the end of induction of iPSC to syndetome-like cells and off-target differentiation to neural-like progenitors. **(A)** UMAPS of each differentiation step sorted into 11 cell clusters from iPSC to SYN were annotated to 6 distinct cell populations: iPSC (OCT4^+^SOX2^+^NANOG+), Syndetome (SYN, MKX^+^TNMD^+^COL1A1^+^), Neuromesodermal Progenitors/Neural Crest cells (NMP/NC, PAX3^+^NRP2^+^COLEC12^+^), Mesoderm (Mes, DLL1^+^DLL3^+^PARAXIS+), Neuromesodermal Progenitors – Cranial (NMP-C, DLL1^+^DLL3^+^NOTCH1^+^CRAB1^+^), and Neural Lineage cells (NL, NRN1^+^DCX^+^NNAT^+^). During induction pluripotent clusters gradually disappeared, and three main clusters emerged: SYN, NMP-C, and NL. (**B-C)** Feature plots **(B)** and dot plots **(C)** of stage-specific genes displayed for all differentiation stages. **(D)** Expression of primary tenogenic markers COL1A1 (blue) and either SCX (red) or TNMD (red) on UMAP plot. **(E-F)** Original samples and clusters were ordered on pseudo-time developmental trajectory. **(E)** Trajectory analysis based on original samples showed transition from iPSC to SYN correlated with the samples, however, SYN cells were located within three endpoints. **(F)** Trajectory analysis based on clusters showed SYN cluster as the main differentiation endpoint with NMP-C and NL as off-target differentiation endpoints. Branching point heat maps **(G-H)** and gene expression were predominated by neural-related markers SYP, PCLO, DCX, and NRN1 **(I)**. **(J-K)** IPA analysis revealed that the off-target clusters NMP-C and NL clusters, were linked with increased Wnt pathway activity **(K-L)**, while SYN cluster was associated with tenocyte differentiation and linked to decreased Wnt pathway activity **(J)**.

### Trajectory analysis shows off-target differentiation to neural-like progenitors

Monocle package (v2.22.0) was used to create pseudo-time trajectory (Fig. 3E-F) and branching point heatmaps (Fig. 3G-H). Examination of branch point heatmaps and marker expression showed that the “side-arm” clusters that predominated in the later SCL stage and especially in SYN (Fig. 3A), were enriched for neural-related markers NRN1, SYP, PCLO and DCX^38–40^ (Fig. 3I). This suggests that these two clusters were likely side products that deviated from the main differentiation trajectory into neural lineage cell population.

To understand the pathways enriched at each induction stage, we used QIAGEN Ingenuity Pathway Analysis (Fig. 3J-L). Examining the canonical pathways in our dataset, we found that in SYN cluster, BMP activation was linked to differentiation towards the tenocyte lineage and concurrent WNT inhibition through DKK1 upregulation (Fig. 3J). Moreover, the side arm clusters showed activation of WNT signaling and upregulation of gene signatures associated with neuronal development (Fig. 3K-L). In conclusion, iPSCs were successfully differentiated into SYN, however, cells expressing neural markers also appeared as side products during the differentiation, and WNT signaling was found to play a role in this process.

### Inhibition of WNT signaling resulted in decrease in off-target differentiation

Pathway analysis showed that the crosstalk between BMP and WNT may play an important role in off-target differentiation products. Thus, we hypothesized that inhibition of WNT signaling may improve induction towards tenogenesis. We investigated the addition of a potent PORCN inhibitor, WNT-C59, which blocks both canonical and non-canonical WNT signaling (Fig. 5A). We observed that when WNT inhibitor (WNTi) was implemented after day 8 (SM stage), it resulted in decreased expression of neural markers NRN1, DYNLLl1 and FMN1 (Fig. 4A). Immunostaining for neural markers (DCX, SYP and NRN1) showed decreased expression in WNTi-treated cells (Fig. 4C1-C3) unlike non-treated cells (Fig. 4B1-B3). Implementation of the WNTi inhibitor also resulted in significantly higher expression of tenogenic marker TNMD partway through SYN induction (Fig. 4A2). A higher proportion of COL1^+^TNMD^+^ and SCX^+^MKX^+^ cells were observed in SYN^WNTi^ compared to SYN group (Fig. 4E-F). Notably, in some cases we observed overgrowth of cells into spherical 3D structures (Fig. 4D), which could be related to the other cell populations comprising ∼50% of this group (Suppl. Table 1 and Suppl. Fig. 2).

**Fig. 4.**
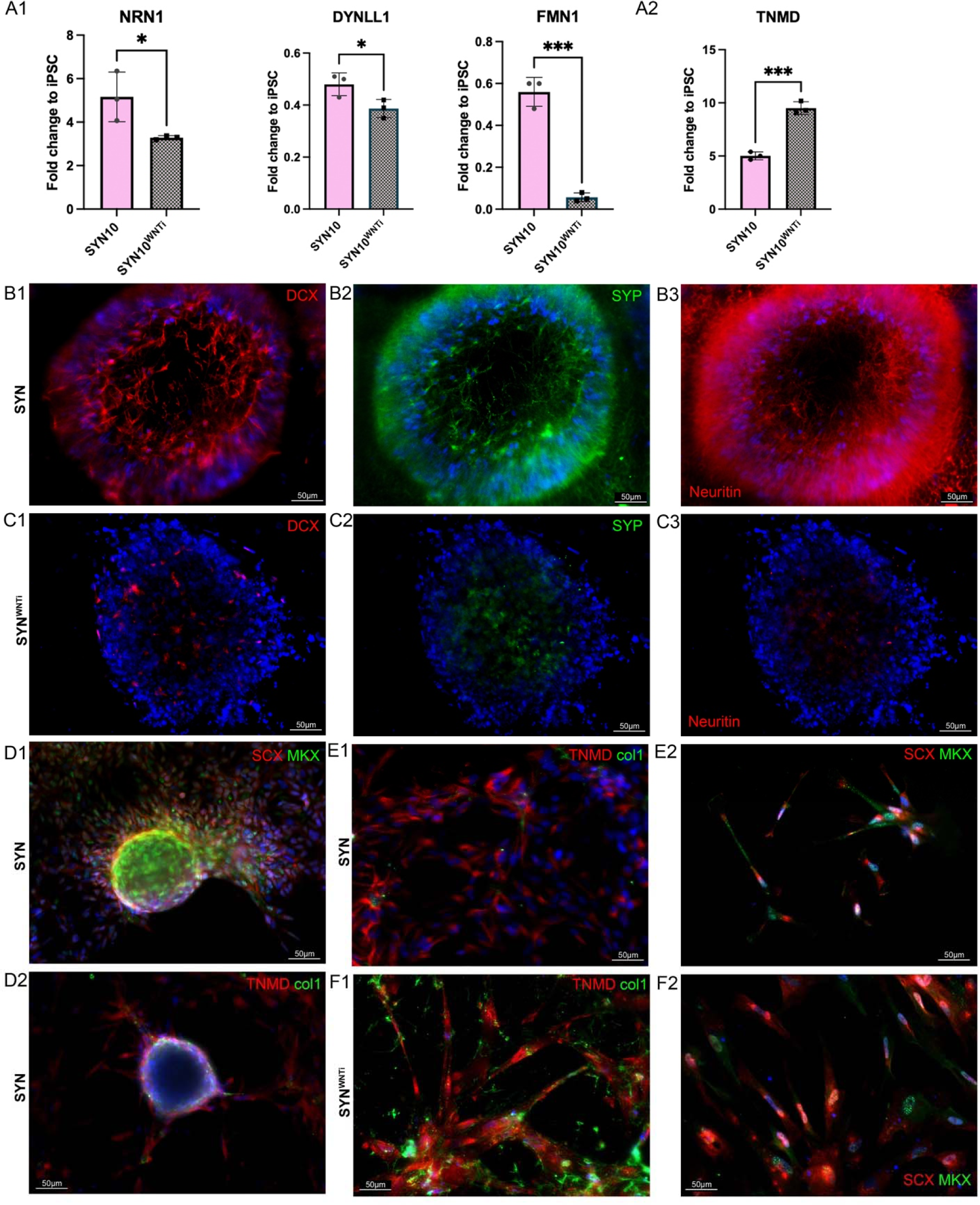
Inhibition of WNT signaling resulted in decreased expression of off-target markers and a more homogeneous population. **(A)** Gene expression analysis of neural markers NRN1, DYNLL1 and FMN1, as well as tenogenic marker TNMD on day 10 of SYN induction (day 21 of the differentiation). **(A1)** Cells treated with WNTi had significantly downregulated neural marker expression compared to just SYN. n=4 replicates/group. **(A2)** Tendon gene expression was significantly upregulated in the WNTi-treated group; n=3/group; *p<0.05, **p<0.01, ***p<0.001, ****p<0.0001. **(B-C)** Immunofluorescence staining for neural markers (DCX, SYP, NRN1), SYN **(B1-B3)** and SYN^WNTi^ **(C1-C3)** showed that they were present in SYN and disappeared in SYN^WNTi^. Scale bars represent 50μm. **(D-F)** Immunofluorescence staining for tenogenic markers (SCX, MKX, TNMD, COL1) of SYN and SYN^WNTi^ showed that more COL1^+^TNMD^+^ and SCX^+^MKX^+^ cells were observed in SYN^WNTi^ compared to SYN. Scale bars are 50μm. Differentiation experiments were repeated independently n=2 with the 007i line and n=2 with a second iPSC line that was later tested (83i). IF staining was performed in n=3 technical triplicates.

**Fig. 5.**
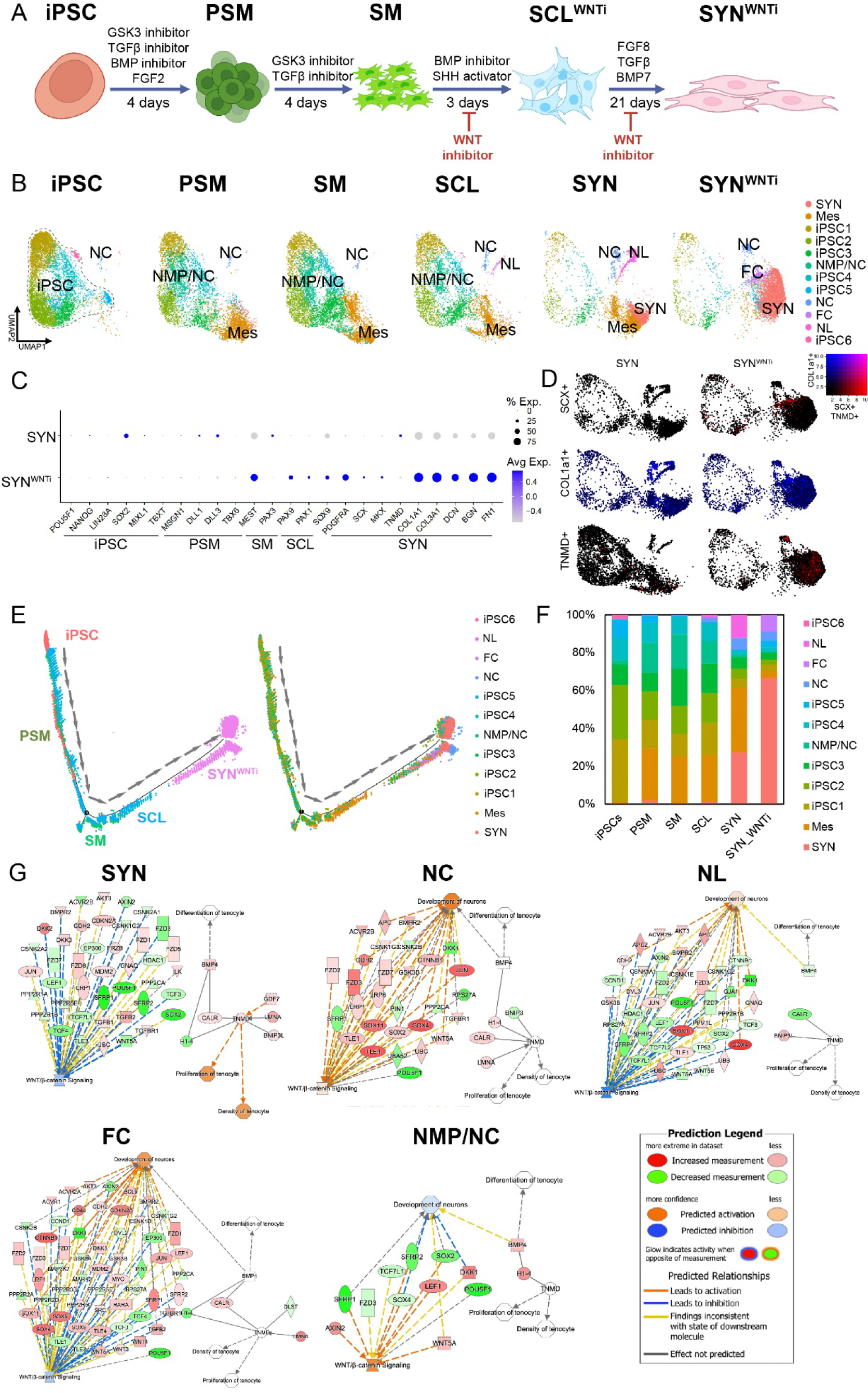
Addition of WNTi to the SCL and SYN induction stages of the differentiation improved differentiation to SYN and eliminated off-target clusters. **(A)** Schematic of optimized iPSC to SYN stepwise induction with WNTi addition implemented at SM to SCL and SCL to SYN stages. Informed by single cell transcriptomics, the addition of Wnt pathway inhibitor at the later stages of the differentiation resulted in a more specific differentiation of iPSCs to tenocytes. **(B)** UMAPS of each differentiation step sorted into 12 distinct clusters from iPSC to SYN were annotated into 6 distinct cell populations: Syndetome (SYN, MKX^+^TNMD^+^DCN^+^BGN^+^), iPSC (OCT4^+^NANOG^+^LIN28A^+^SOX2^+^), Neuromesodermal Progenitors/Neural Crest (NMP/NC, TBXT^+^TWIST1^+^SP5^+^SNAI2^+^), Neural Crest (NC, PTN^+^NTKR2^+^SOX4^+^SOX11^+^), Fibrocartilage (FC, COL2A1^+^SOX9^+^FN1^+^BGN^+^COL1a1^+^), Neural Lineage (NL, SOX2^+^DCX^+^MAP2^+^UNCX^+^SOX4^+^). UMAP comparison of the SYN and SYN^WNTi^ populations demonstrated increased size of the SYN cluster and elimination of NL cluster. **(C)** Dot plots of gene expression of stage-specific genes for SYN and SYN^WNTi^. **(D)** Expression of primary tenogenic markers Col1a1 (blue), Scx (red) and Tnmd (red) on UMAP plot for SYN and SYN^WNTi^. **(E)** Original samples and clusters were ordered on pseudo-time developmental trajectory. Trajectory analysis revealed one main endpoint, the SYN cell populations. **(F)** After addition of WNTi, the proportion of cells in the SYN clusters increased by 59% while the proportion of cells in the NL cluster was eliminated. **(G)** IPA network analysis showed that WNT pathway was enriched in the NMP/NC and NC clusters but not in NL.

To further unveil the role of continuous WNT inhibition at the SM stage and until the end of the induction to SYN, we performed scRNA-seq (Fig. 5A). 5645 cells were subjected to unsupervised clustering and plotted with UMAPs. Integrated UMAP plots of all the cell populations in this study revealed 12 distinct clusters which were further characterized. We annotated 7 cell populations by using classical developmental markers and performing gene ontology enrichment analysis for DEGs (Fig. 5B). Six clusters expressed pluripotency and stem cell markers, including NANOG, OCT4, and SOX2 and were annotated as pluripotent cells (iPSC1-6). These populations were evident during the first stages of induction, and they gradually receded. At PSM stage two mesodermal populations were identified, one was annotated as Mes (TBXT^+^DLL1^+^) and another as NMP-C, where TWIST1,^25^ SP5,^41^ and SNAI2^42^ were upregulated. These two populations further expanded at the SM stage. At the SCL stage, two additional newly formed clusters appeared. These were categorized as NL and neural crest (NC) cells. The NL cells expressed markers associated with neurogenesis including CRABP1, DCX, MAP2.^43^ The NC cells expressed SOX4, SOX11, and NCAM1.^44,45^ At the SYN stage we identified a SYN population with cells expressing tendon and ECM associated markers (COL1A1^+^, COL1A2^+^, COL3A1^+^, COL11A1^+^, FN1^+^, FBN1^+^, DCN^+^). WNT inhibition resulted in the elimination of the NC population, while the SYN population has increased in size and a second population expressing tendon and some chondrogenic markers was evident, denoted as fibrocartilage (FC) cluster. The expression levels of classical stage-specific markers in SYN^WNTi^ presented in a dot plot showed increased expression compared to untreated SYN group (Fig. 5C). Density plots of cells expressing COL1A1 and tendon markers SCX and TNMD showed increased positive cells for those markers in SYN^WNTi^ compared to SYN group (Fig. 5D). Trajectory analysis for the WNTi-treated group showed elimination of the bifurcations that were observed in the non-treated group (Fig. 5E). Cell proportions for each cluster were calculated by normalizing to total cell numbers. It was shown that 75.6% of all cells in SYN^WNTi^ were clustered at the two SYN subpopulations, SYN and FC (66.9 and 8.7%, respectively; Fig. 5F). Comparison of WNTi treated groups using a second iPSC line (83i) showed that in the second line, 78% of cells were clustered in the SYN group, while the FC cluster was non-existent (Suppl. Fig. 3 and Suppl. Table 1).

IPA network analysis following treatment with the WNTi showed that WNT pathway activation is associated with the NMP-C and NC clusters and not with the SYN, NL, and FC clusters (Fig. 5G).

## Discussion

In this study, informed by ontogenesis and recent reports modeling PSC differentiation *in vitro*, we successfully differentiated human iPSCs to syndetome-like cells in a stepwise manner using a GMP-ready line and chemically defined serum-free media through balancing the BMP, TGFβ Activin/Nodal, FGF, and WNT signaling pathways. iPSCs were differentiated to presomitic mesoderm, somite, sclerotome and syndetome-like cells (Fig.1A-B). Gene expression analysis, IF, and scRNA-seq analysis validated the stepwise differentiation using classical markers that define each developmental step. However, gene regulatory network analysis revealed that the major off-target cell populations were highly associated with WNT signaling activation beginning from the SCL stage and into the SYN stage. Using scRNA-seq, we confirmed our hypothesis that WNT inhibition with the WNT signaling inhibitor (WNTi) Wnt-C59 blocked cell fate bifurcation to the “unwanted” neural fates and drove induction towards tenogenesis.

Gene expression analysis showed significant upregulation of stage-specific markers (Fig. 1C-G). PSM induction was assessed via the Notch receptor ligand Delta-like protein-1 (DLL1) gene and protein expression. Somites form from the epithelization and segmentation of paraxial mesoderm along the rostro-caudal (RC) axis of the embryo in the process of Mesenchymal to Epithelial Transition (MET). Studies in several animal models have shown that that somite RC polarity is established prior to SM formation and it is orchestrated by NOTCH signaling.^24^ Many DLL-family proteins, which are NOTCH receptor ligands, are central to this process, with DLL1 shown as the most critical during PSM formation.^35,46^ Here, we found that the initial seeding density of iPSCs impacted DLL1 expression and thus induction efficiency. Higher starting densities resulted in decreased DLL1^+^ cell ratios assessed by flow cytometry (Suppl. Fig. 1). This is in line with several reports that have identified starting PSC seeding density as an additional factor influencing cell differentiation in animal models and human cells.^47^ This could be attributed to auto/paracrine factors and cell-cell contact.^48^ Unfortunately, iPSC seeding density is often not reported in published protocols and it may be contributing to discrepancies and reproducibility concerns between different laboratories. Taken together, our data indicate that starting seeding density is a crucial component of chemical induction which could influence reproducibility and translatability of a given approach.

Somitogenesis induction was achieved by TGFβ inhibition (SB431542) and moderate WNT activation (CHIR99021) similar to previous reports.^26^ Combined SHH signaling activation (SAG) and BMP inhibition (LDN193189) resulted in induction of SM to SCL. Last, stepwise FGF8 followed by BMP and TGFβ administration induced SYN formation. Early tendon-associated markers were significantly upregulated at the final timepoint of SYN induction (Fig. 1G). FGF ligands including FGF8 are secreted by the myotome region of the somite, located next to the sclerotome, inducing Scleraxis (SCX) expression, which in turn drives syndetome specification.^24^ Protein expression at each stage was verified with IF (Fig. 2). However, at the end of differentiation we noticed gradual cell attrition in the cultures (Fig. 1B, 2I). This could be attributed to the heterogeneity of the cell population at SCL stage, which was confirmed by scRNA-seq (Fig. 3A).

In the current study, scRNA-seq analysis showed that mesodermal cells persisted until SYN induction (Fig. 3A-C). DLL1 expression was significantly upregulated at PSM and downregulated at SM stage. At PSM, 96.2% of the cell population was DLL1^+^ (Suppl. Fig. 1). However, dot plots showed some DLL1 and DLL3 expression in all stages and clusters (Fig. 3C and Suppl. Fig. 4). Consequently, using a single marker may not be sufficient to assess lineage differentiation efficiency. Trajectory analysis showed two distinctive bifurcations at SM and SCL stages. The first branchpoint was observed at the SM stage, when a Mes (DLL1^+^DLL3^+^PARAXIS^+^) and a transient NMP (PAX3^+^NRP2^+^COLEC12^+^) cell population were identified. NMPs were present at SM and disappeared thereafter, while at SCL two “side-arm” clusters evolved (Fig. 3A). The NL cluster was enriched for several neural-associated markers (NRN1^+^NNAT^+^CRABP1^+^ONECUT2^+^NPTX2^+^). In the NMP-C side-arm cluster the top DEGs were a combination of neural-associated markers (NOTCH1^+^CRABP1^+^GREB1^+^ID4^+^), cranial mesodermal markers (DLL1^+^DLL3^+^) and ECM genes including collagens and FN1. The emergence of these two off-target cell populations became quite prominent at SCL, while they were further amplified at SYN (Fig. 3A).

Sclerotome is derived from the ventromedial part of somites and its induction is orchestrated by SHH.^24^ It is established that SHH and NOG produced in the notochord and neural tube stimulate MET, which produces both NC cells from the ectoderm as well as SCL at the ventromedial aspect of somites.^49^ Different groups have achieved sclerotome induction of iPSCs with varying approaches. For instance, Loh *et al* identified that WNT and SHH acted antagonistically in SCL specification.^25^ The NMP-C off-target cell population that was first observed in the SCL and SYN groups could have arisen from NMPs (Fig. 3A). NC, which are a transient cell population of ectodermal origin, are derived from bipotential NMPs from the primitive streak in the presence of FGF, WNT and likely medium BMP activity.^50,51^ However, the timing and levels of BMP signaling required for NC differentiation are still not well understood. Wu *et al*^12^ observed NC off-target populations during stepwise iPSC to somite to chondroprogenitor differentiation. In that study, the authors first observed NC at the chondroprogenitor stage, and hypothesized that they emerged due to BMP treatment for 3 days at the SCL stage.^12^ In the present study, we observed the emergence of NC cells earlier and even though we applied a BMP inhibitor during the SM to SCL differentiation, it may not have been sufficient to modulate BMP activation. Lastly, the NC off-target population could have been derived from NMP/NC cells or from NMPs present at SM. *In vivo* somites give rise to the dermatome and myotome dorsally and the SCL ventromedially.^24^ The dermatome sits next to the ectoderm and can differentiate into the meningotome which can form spinal meninges as well as fibroblasts of the meninges and arachnoids.^52^ The off-target neural cluster in the present study showed increased expression of markers associated with NC as well as neural markers. Thus, it could be posited that it was derived from the NC off-target subpopulation identified at SCL.

Next, DEG analysis using the IPA platform revealed the interaction between WNT and BMP signaling at the cornerstone between the SYN cluster and the NL and NC off-target “side-arm” clusters (Fig. 3J-L). More specifically, upregulation of genes associated with WNT signaling activation in both off-target clusters were observed. Therefore, a WNT pathway inhibitor was employed during SCL and SYN differentiation, which resulted in significant downregulation of selected neural markers, assessed by RT-qPCR and IF (Fig. 4).

Treatment with the WNTi following the SM stage resulted in the expansion of the SYN clusters (Fig. 5A) as well as complete elimination of the NL off-target cluster (Fig. 5A). Further, a new cell population was observed only in the WNTi-treated group, expressing some chondrogenic markers (COL2A1^+^SOX9^+^FN1^+^BGN^+^COL1A1^+^), tenogenic genes and Klf transcription factors, which were recently shown to be associated with bipotential tendon-to-bone attachment cells.^53^ In the same study, these cells were shown to activate a combination of tendon and chondrogenic transcriptomes. Fibrocartilage is found in the enthesis of tendons and shares the same progenitors as the syndetome.^54,55^ Intriguingly, the FC cluster was not observed in the second cell line we differentiated (Suppl. Fig. 3) and it was the only observed difference between the two lines. Fibrocartilage progenitors express SCX, however, divergence of SCX^+^ cells towards tendon or fibrocartilage have yet to be fully understood. In this work, WNT inhibition at the SM stage resulted in increased numbers of SCX^+^ derivatives compared to SYN while eliminating the NL population. However, the NC cell population was not diminished, indicating that these cells might have been generated due to “contaminating” multipotential NMPs still present at SM and SCL. IPA network analysis revealed WNT pathway activation in mesodermal and NC clusters and not the final products of the induction, that is NL, FC, and SYN clusters (Fig. 5G). This further confirms our hypothesis regarding its activation following SM specification and suggests that it plays an important role in specifying the fate of NMPs towards neural lineages. Further studies are needed to delineate the level of BMP activity required to either induce or block bifurcation of NMPs to NC fates. Interestingly, SCX expression peaked in the middle of SYN induction and decreased at the end (Suppl. Fig. 5), when TNMD, COLA1A and other tenogenic markers (collagens, DCN, BGN1, FN1) were upregulated. This suggests that SYN induction could be further optimized and shortened. Further, WNT inhibition during the entirety of the SYN induction might potentially negatively impact SYN maturation and it should be further fine-tuned. For instance, WNT signaling has been shown to upregulate TNMD expression in equine BM-MSCs,^56^ while it was shown to suppress TNMD and other tenogenic genes in tendon-derived cells *in vitro.*^57^

Previous studies investigating iPSC to SYN differentiation reported varying yields at the end of induction. Nakajima *et al* reported 68% SCX+ cells using flow cytometry.^26^ In a follow up study, the same research group reported 91.6% SCX+, 90.4% MKX+, 79.9% COL1A1+ and 77.5% COL1A2+ protein expression using IF quantification.^14^ However, although informative, it could be noted that IF is only a semi-quantitative assessment burdened with operator bias and lower sensitivity compared to flow cytometry or scRNA-seq, unless performed in a more automated manner.^58^ Nevertheless, despite those caveats, the high individual expression of all four markers (77.5%-91.6%) with IF supports a relatively efficient SYN differentiation overall. Kaji *et al* differentiated mESCs from SCX-GFP reporter mice to SYN and reported 90% efficiency assessed by flow cytometry for SCX,^15^ while Sugimoto *et al* showed 66% efficiency utilizing a similar Scx-GFP mouse model combined with flow cytometry and scRNA-seq assessments.^15,16^ In the current study, SYN clusters, characterized by scRNA-seq analyses, greatly increased in the SYN^WNTi^ group, from 47.6% prior to WNTi application, to 67%-78% of total cells after treatment (Fig. 5F and Suppl. Fig. 3). In conclusion, we showed that WNT inhibition was able to successfully modulate the later stages of differentiation from SM to SYN.

This study is not without limitations. The IPA network analysis is a knowledge-based and hypothesis driven platform. We have specifically targeted known pathways to be involved in syndetome differentiation. However, WNT signaling stood out with very specific affinity to the off-target populations and we have verified our findings with experiments proving this hypothesis. The SCL to SYN induction, maturation, and expansion in culture was considerably long and resulted in decreased expression of SCX (Suppl. Fig. 5), suggesting that it could be shortened. Even though following WNTi treatment induction efficiency increased considerably, there were still other populations present, including NC and undifferentiated cells. Cell sorting could be used to isolate a more homogeneous cell population. Further, removal of undifferentiated cells is crucial for *in vivo* preclinical studies. Future work is warranted to assess the functionality of the cells *in vivo*.

## Conclusions

Tendons have poor innate healing capacity and novel approaches are urgently needed. Tendon cell therapy offers potential for regeneration of injured tissues. In this study, we successfully differentiated a GMP-ready human iPSC line to syndetome-like cells in a stepwise manner using fully defined media. However, scRNA-seq trajectory analysis revealed off-target differentiation towards a neural phenotype, resulting in heterogeneous differentiation. Differentially expressed gene and regulatory network analyses demonstrated WNT-associated effectors implicated in the off-target groups, which prompted us to apply a WNT inhibitor at the somite stage. Taken together, our data provide evidence that by manipulating WNT signaling we can achieve a more specific and robust differentiation of iPSCs to tendon progenitors. Elucidating the mechanism of the WNT signaling pathway using a development-inspired protocol can lead to the development of more powerful and specific differentiation protocols for cell therapy applications. iPSC-derived tendon progenitors can be an off-the-shelf cell source to tendon and ligament injuries that are currently untreatable or have poor surgical outcomes.

## Acknowledgements

This study was supported by Cedars-Sinai Board of Governors Regenerative Medicine Institute, the NIH/NIAMS K01AR071512 (DS), California Institute for Regenerative Medicine DISC0-14350 and EDUC4-12751 (WJ) awards.

## Author contributions

Conceptualization, AP and DS; Methodology, AP and DS; Software, WJ and JS; Validation, AP, VY, WJ and JS; Formal analysis, AP, VY and WJ; Investigation, AP, VY, TS, AC, CC and MC; Resources, AP, VY and JS; Data Curation, WJ and JS; Writing-Original Draft, AP, VY and DS; Visualization, VY, AP and JS; Funding Acquisition, DS and WJ.

## STAR Methods

### Resource availability

#### Lead contact

Further information and requests for resources, reagents, data, and code should be directed to the corresponding author, Dmitriy Sheyn.

#### Materials availability

This study did not produce any unique reagents or materials.

#### Data and code availability

The raw sequencing data and original data are accessible in Gene Expression Omnibus. The series accession number is GSE229008. The original code is accessible at GitHub: https://github.com/jason199112345/scRNA-seq-for-the-project-of-iPSC-to-Tenocyte-differentiation.git.

### Experimental model and subject details

#### iPSC maintenance and expansion

Human iPSC lines were obtained from the Cedars-Sinai Core facility and were expanded on Matrigel^TM^-coated plates (BD Biosciences) (0.08 mg/well in 6 well plates). Cells were fed every other day with cGMP mTeSR™Plus media (StemCell Technologies) and passaged with ReLeSR™ (StemCell Technologies). Two distinct fully characterized hiPSC lines were used in this study: the GMP-ready CS0007iCTR-n5 line and the CS83iCTR-22n1 line (https://biomanufacturing.cedars-sinai.org/?filter_cell-type=ipsc&filter_primary-tissue&filter_disease&filter_sex&filter_age-at-sampling&filter_ethnicity&filter_race&filter_gene&filter_mutation&filter_project&cs_product_search).

## Method details

### iPSC to SYN differentiation

iPSCs were seeded into Matrigel-coated plates (#354230, Corning, Corning, NY) (0.08 mg/well in 6 well plates and 0.04mg/well in 12 well plates) and were induced to presomitic mesoderm (PSM) at 30% confluency (∼50-60 aggregates/cm^2^) in basal medium (BM) composed of IMDM/Ham’s F12 (1:1) (Thermo Fisher), 1% lipid concentrate (Thermo Fisher), 0.5% antibiotic-antimycotic solution (Thermo Fisher), 15 µg/mL apo-transferrin (Sigma), 450µM monothioglycerol (Sigma), 7µg/mL insulin (Sigma) supplemented with 10 µM SB431542 (Sigma), 2µM DMH1 (Santa Cruz Biotechnology), 20ng/mL FGF2 (Biogems-PeproTech), and 10µM CHIR99021 (Biogems-PeproTech) (PSM media). After 3 days, media were changed to fresh PSM media. Geltrex (#A1413301, Thermo Fisher) was diluted 1:100 in cold serum-free DMEM/F12 to coat wells in 12-well plates (0.5mL/well). The coated plates were incubated for at least 1hr at 37°C before use. At day 4, PSM cells were washed with PBS and lifted using Accutase (StemCell Technologies) for 5min at 37°C and reseeded at 28,500 cells/cm^2^ into the Geltrex-coated plates supplemented with SM differentiation media composed of BM supplemented with 10µM SB431542, 5µM CHIR99021, and 10µM Y-27632 dihydrochloride (Biogems, CA, USA). On days 5 and 7, media were changed to fresh SM media without Y-27632 dihydrochloride. At day 8, cells were cultured in sclerotome (SCL) differentiation media containing BM supplemented with 100nM SAG (Biogems-PeproTech), and 0.6µM LDN193189 (Biogems-PeproTech). Media were changed to fresh SCL media at day 10. At day 11, cells were washed with PBS, lifted using Accutase (StemCell Technologies) for 5min at 37°C and were reseeded at 60,000 cells/cm^2^ into Geltrex-coated (Thermo Fisher) plates with BM supplemented with 20ng/mL FGF8 (Peprotech). After 3 days, media were changed to BM supplemented with 10ng/mL TGFβ3 (StemCell Technologies), and 10ng/mL BMP7 (PeproTech) (SYN maturation, SYN-M media). Media were changed to fresh SYN-M media every 3 or 4 days until day 32.

### Application of Wnt inhibitor

Following initial single cell analyses, the iPSC to SYN induction was repeated with the addition of the Wnt signaling inhibitor Wnt-C59 (Cayman Chemical, MI, USA) beginning from the SCL induction stage and onwards; the somite cells treated with SCL media and sclerotome cells treated with SYN-M media were supplemented with 1µM Wnt-C59.

### Flow cytometry

For assessment of DLL-1 levels to determine induction efficiency, cells that were induced for 4 days were lifted using Accutase for 5min at 37°C and washed 3x with FACS buffer containing 2% bovine serum albumin (BSA, A4503, Sigma, St. Louis, MO) and 0.1% sodium azide (S2002, Sigma) in PBS. The cells were either unstained, labeled with DLL1 anti-human antibody (APC-Vio 770, #130-106-148, Miltenyi Biotec, Bergisch Gladbach), or labeled with its associated isotype control (APC-Vio 770, #130-113-759, Miltenyi Biotec) for 15min protected from light at 4°C. After the incubation, cells were washed again and resuspended with FACS buffer. Data was acquired on a BD LSR Fortessa analyzer (BD Biosciences, San Jose, CA) and was analyzed using FlowJo software (FlowJo LLC, Ashland, OR).

### Gene expression analysis

Differentiation to SYN was defined based on expression of developmental stage-specific markers (Supplemental Table 1). Cells were collected from each developmental stage and total RNA was isolated using the RNeasy plus kit (Qiagen), and reverse transcribed with the high-capacity cDNA reverse transcription kit (Applied Biosystems). Then, cDNA was amplified by performing qPCR with TaqMan^®^ gene expression. The threshold cycle (Ct) value of 18S rRNA was used as an internal control using the TaqMan^®^ gene expression FAM/MGB probe system (4333760F, Thermofisher). The Livak method was used to calculate ΔΔCt values and fold change was calculated as 2^-ΔΔCt^, as previously described and published.^59^

### Immunocytochemistry for phenotype confirmation

In preparation for immunocytochemistry, iPSCs were seeded onto coverslips. The coverslips were coated with Cultrex Poly-D-Lysine (R&D Systems) for 5min at RT, aspirated, and air-dried for at least 2h. They were then coated with Matrigel as per the seeding protocol described previously. At each developmental stage, cells were fixed with 4% paraformaldehyde for 30min at RT, followed by 3x PBS washes. Briefly, after serum-free protein blocking (X0909, Dako, Agilent), cells were hybridized with various primary antibodies (Table 1) overnight at 4°C. The following day, these cells were incubated with fluorophore-conjugated secondary antibodies (1h at 37°C). Coverslips were mounted onto slides using ProLong® Gold with DAPI (Molecular Probes, Life Technologies). A Carl Zeiss fluorescence microscope (Imager, Z1, ApoTome and MBF equipped) was used to acquire images.

### Single cell RNA sequencing sample preparation, sequencing, and data processing

At each developmental stage (iPSC, PSM, SM, SCL, SYN, SYN^WNTi^), cells were washed with PBS and lifted using Accutase (StemCell Technologies) for 5min at 37°C, and then centrifuged for 3min at 1000RPM. The cells were then resuspended in media, filtered using a 40 µM FlowmiTM cell strainer (Thermo Fisher) and washed twice more with media before being filtered for the last time to ensure a single cell suspension. The filtered cells were manually counted in quadruplicate with 0.4% trypan blue dye (Thermo Fisher) and cells were resuspended in media at a concentration of 1,500 cells/µL.

Single-cell RNA-Seq libraries were prepared per the Single Cell 3′ v3.1 Reagent Kits User Guide (10x Genomics, Pleasanton, California) using the 10x Genomics Chromium Controller. Barcoded sequencing libraries were quantified by quantitative PCR using the Collibri Library Quantification Kit (Thermo Fisher Scientific, Waltham, MA). Libraries were sequenced on a NovaSeq 6000 (Illumina, San Diego, CA) as per the Single Cell 3′ v3.1 Reagent Kits User Guide, with a sequencing depth of ∼40,000 reads/cell.

Raw sequencing data was demultiplexed and converted to FASTQ format by using bcl2fastq v2.20 (Illumina, San Diego, California). More than 200 million reads were obtained for each sample. Reads were mapped to the human GRCh38 genome and count quantification was done using 10x Genomics Cell Ranger v.7.0.0.

We utilized our previously established single-cell analysis platform based on R (v4.1.2) ^60^. Seurat package (v4.1.0) was used to load and process the single-cell data. Specifically, the data for each sample were loaded using *Read10X*. Quality control was performed for each sample. The parameters used for quality control were *nFeature_RNA*, that is the counts of genes detected in each cell, and *percent.mt*, that is the percent of mitochondrial genes expressed in each cell. Any cells having *nFeature_RNA* outside of the range of 200-8000 were excluded in the downstream analysis. Any cells with the *percent.mt* being smaller than 20%, were excluded in the downstream analysis. The Seurat object for each sample after the quality control was combined into a larger Seurat object. The large Seurat object was then normalized, anchored, and integrated. Principal component analysis with PC=30 was performed, dimensional reduction with Uniform Mani-fold Approximation and Projection (UMAP) was applied, and clusters were identified in an unsupervised manner with a resolution of 0.5. Following the identification of clusters, each cell was annotated with two important identities: the sample they were derived from, and the cluster (cell type) they belonged to. The differentially expressed genes (DEG) were calculated by comparing across sample identity or cluster identity, i.e., a specific cluster vs. all cells, a specific sample vs. all other samples, a specific sample vs. another specific sample, or as otherwise noted in the manuscript. The DEGs were inputted to Qiagen Ingenuity Pathway Analysis (IPA) for gene ontology (GO) term enrichment and pathway analysis. Genes with p<0.0001 were included in the Qiagen IPA analysis.

The pseudo-time trajectory was constructed using the Monocle package (v2.18.0). The previously combined large Seurat object was loaded into the Monocle package. DDRTree method was used to reduce the dimension of the data. The cells were projected on the trajectory and visualized based on their identity or marker expression.

### Statistical analyses

Data are presented as mean ± standard deviation from the mean. Normally distributed data were analyzed with unpaired t-test (for 2 groups), or non-repeated measures analysis of variance followed by Tukey-Kramer HSD *post hoc* analysis when more than 2 groups were compared. Non-parametric data were analyzed using the Mann-Whitney and Kruskal-Wallis tests. Statistical significance was set at p<0.05.

## Supplemental information titles and legends

**Suppl. Table 1:** Normalized cell counts (expressed as %) per cluster following WNTi treatment, shown for the two cell lines, 007i and 83i.

**Suppl. Fig 1: Flow cytometry of DLL-1 at the PSM stage for low and high seeding densities. Left panel:** low iPSC seeding density resulted in high percentage of DLL-1+ cells. **Right panel**: high iPSC seeding density resulted in a comparably reduced DLL-1+ cell population.

**Suppl. Fig 2. Distribution of cell subpopulations per sample.** Cell clusters from iPSC to SYN were annotated to 6 distinct cell populations: iPSC (OCT4^+^SOX2^+^NANOG^+^), Syndetome (SYN, MKX^+^TNMD^+^COL1A1^+^), Neuromesodermal Progenitors/Neural Crest (NMP/NC, PAX3^+^NRP2^+^COLEC12^+^), Mesoderm (Mes, DLL1^+^DLL3^+^PARAXIS^+^), Neuromesodermal Progenitors – Cranial (NMP-C, DLL1^+^DLL3^+^NOTCH1^+^CRAB1^+^), and Neural Lineage cells (NL, NRN1^+^DCX^+^NNAT^+^).

**Suppl. Fig 3. *A).*** ScRNA-seq comparison of two different iPSC lines for SYN^WNTi^. Cell population annotation is shown in Fig.5 and in Suppl. Table 1. ***B).*** Distribution of cell subpopulations per sample, where the second cell line sample at the syndetome stage has also been included (designated as WNTi-83i). Clusters numbers and annotations are shown in Suppl. Table 1.

**Suppl. Fig. 4**. **Dot plot of stage-specific markers of SYN^WNTi^ for each cluster.** Cell population annotation is presented in Fig. 5.

**Suppl. Fig. 5. Change in SCX expression throughout SYN induction with WNTi.** All timepoints are normalized to day 0 (iPSC). *p<0.0001.

**Suppl. Fig. 6.** A non-biased GO analysis was performed. Multiple pathways were detected in the three pathways of interest, that is C3 (Fig. 3. A; SYN), C9 (Fig. 3A; NMP/NC) and C10 (Fig. 3A ;NL).

**Suppl. Fig. 7.** Immunofluorescence staining of selected markers and DAPI for nuclear staining from Fig.2 expanded to show all separate channels. (B’) NANOG at PSM stage; (E’), MEOX1 and PAX3 at SCL stage (G’) PAX9 at SM stage.

**Suppl. Fig. 8.** Separate and combined feature plots of SCX and TNMD expression at SCL through SYN stage with and without WNT inhibitor (WNTi).

**Suppl. Fig. 9.** Feature plots of NKX3.2 and MEOX1 displayed for all differentiation stages.

## References

1. Yang, G., Rothrauff, B.B., and Tuan, R.S. (2013). Tendon and ligament regeneration and repair: clinical relevance and developmental paradigm. Birth Defects Res C Embryo Today 99, 203–222. 10.1002/bdrc.21041.

2. Liu, W., Yin, L., Yan, X., Cui, J., Liu, W., Rao, Y., Sun, M., Wei, Q., and Chen, F. (2017). Directing the Differentiation of Parthenogenetic Stem Cells into Tenocytes for Tissue-Engineered Tendon Regeneration. Stem Cells Transl Med 6, 196–208. 10.5966/sctm.2015-0334.

3. Webster, K.E., Feller, J.A., Leigh, W.B., and Richmond, A.K. (2014). Younger patients are at increased risk for graft rupture and contralateral injury after anterior cruciate ligament reconstruction. Am J Sports Med 42, 641–647. 10.1177/0363546513517540.

4. Puetzer, J.L., Ma, T., Sallent, I., Gelmi, A., and Stevens, M.M. (2021). Driving hierarchical collagen fiber formation for functional tendon, ligament, and meniscus replacement. Biomaterials 269, 120527.

5. Cai, J., Xie, X., Li, D., Wang, L., Jiang, J., Mo, X., and Zhao, J. (2020). A novel knitted scaffold made of microfiber/nanofiber core–sheath yarns for tendon tissue engineering. Biomaterials Science 8, 4413–4425.

6. Ho, J.O., Sawadkar, P., and Mudera, V. (2014). A review on the use of cell therapy in the treatment of tendon disease and injuries. Journal of Tissue Engineering 5, 2041731414549678.

7. Abbah, S.A., Spanoudes, K., O’Brien, T., Pandit, A., and Zeugolis, D.I. (2014). Assessment of stem cell carriers for tendon tissue engineering in pre-clinical models. Stem cell research & therapy 5, 1–9.

8. Jo, C.H., Lim, H.J., and Yoon, K.S. (2019). Characterization of Tendon-Specific Markers in Various Human Tissues, Tenocytes and Mesenchymal Stem Cells. Tissue Eng Regen Med 16, 151–159. 10.1007/s13770-019-00182-2.

9. Wang, S., Bates, J., Li, X., Schanz, S., Chandler-Militello, D., Levine, C., Maherali, N., Studer, L., Hochedlinger, K., Windrem, M., and Goldman, Steven A. (2013). Human iPSC-Derived Oligodendrocyte Progenitor Cells Can Myelinate and Rescue a Mouse Model of Congenital Hypomyelination. Cell Stem Cell 12, 252–264. 10.1016/j.stem.2012.12.002.

10. Kanke, K., Masaki, H., Saito, T., Komiyama, Y., Hojo, H., Nakauchi, H., Lichtler, Alexander C., Takato, T., Chung, U.-i., and Ohba, S. (2014). Stepwise Differentiation of Pluripotent Stem Cells into Osteoblasts Using Four Small Molecules under Serum-free and Feeder-free Conditions. Stem Cell Reports 2, 751–760. 10.1016/j.stemcr.2014.04.016.

11. Yamashita, A., Morioka, M., Yahara, Y., Okada, M., Kobayashi, T., Kuriyama, S., Matsuda, S., and Tsumaki, N. (2015). Generation of Scaffoldless Hyaline Cartilaginous Tissue from Human iPSCs. Stem Cell Reports 4, 404–418. 10.1016/j.stemcr.2015.01.016.

12. Wu, C.-L., Dicks, A., Steward, N., Tang, R., Katz, D.B., Choi, Y.-R., and Guilak, F. (2021). Single cell transcriptomic analysis of human pluripotent stem cell chondrogenesis. Nature communications 12, 362.

13. Komura, S., Satake, T., Goto, A., Aoki, H., Shibata, H., Ito, K., Hirakawa, A., Yamada, Y., and Akiyama, H. (2020). Induced pluripotent stem cell-derived tenocyte-like cells promote the regeneration of injured tendons in mice. Scientific reports 10, 3992.

14. Nakajima, T., and Ikeya, M. (2021). Development of pluripotent stem cell-based human tenocytes. Development, Growth & Differentiation 63, 38–46.

15. Kaji, D.A., Montero, A.M., Patel, R., and Huang, A.H. (2021). Transcriptional profiling of mESC-derived tendon and fibrocartilage cell fate switch. Nature Communications 12, 4208.

16. Yoshimoto, Y., Uezumi, A., Ikemoto-Uezumi, M., Tanaka, K., Yu, X., Kurosawa, T., Yambe, S., Maehara, K., Ohkawa, Y., and Sotomaru, Y. (2022). Tenogenic Induction From Induced Pluripotent Stem Cells Unveils the Trajectory Towards Tenocyte Differentiation. Frontiers in Cell and Developmental Biology 10, 780038.

17. Donderwinkel, I., Tuan, R.S., Cameron, N.R., and Frith, J.E. (2022). Tendon tissue engineering: Current progress towards an optimized tenogenic differentiation protocol for human stem cells. Acta Biomaterialia.

18. Brown, J.P., Galassi, T.V., Stoppato, M., Schiele, N.R., and Kuo, C.K. (2015). Comparative analysis of mesenchymal stem cell and embryonic tendon progenitor cell response to embryonic tendon biochemical and mechanical factors. Stem cell research & therapy 6, 1–8.

19. Havis, E., Bonnin, M.-A., Olivera-Martinez, I., Nazaret, N., Ruggiu, M., Weibel, J., Durand, C., Guerquin, M.-J., Bonod-Bidaud, C., and Ruggiero, F. (2014). Transcriptomic analysis of mouse limb tendon cells during development. Development 141, 3683–3696.

20. Havis, E., Bonnin, M.-A., Esteves de Lima, J., Charvet, B., Milet, C., and Duprez, D. (2016). TGFβ and FGF promote tendon progenitor fate and act downstream of muscle contraction to regulate tendon differentiation during chick limb development. Development 143, 3839–3851.

21. Dale, T.P., Mazher, S., Webb, W.R., Zhou, J., Maffulli, N., Chen, G.-Q., El Haj, A.J., and Forsyth, N.R. (2018). Tenogenic differentiation of human embryonic stem cells. Tissue Engineering Part A 24, 361–368.

22. Schweitzer, R., Zelzer, E., and Volk, T. (2010). Connecting muscles to tendons: tendons and musculoskeletal development in flies and vertebrates. Development 137, 2807–2817.

23. Huang, A.H., Lu, H.H., and Schweitzer, R. (2015). Molecular regulation of tendon cell fate during development. Journal of Orthopaedic Research 33, 800–812.

24. Tani, S., Chung, U.-i., Ohba, S., and Hojo, H. (2020). Understanding paraxial mesoderm development and sclerotome specification for skeletal repair. Experimental & Molecular Medicine 52, 1166–1177.

25. Loh, K.M., Chen, A., Koh, P.W., Deng, T.Z., Sinha, R., Tsai, J.M., Barkal, A.A., Shen, K.Y., Jain, R., and Morganti, R.M. (2016). Mapping the pairwise choices leading from pluripotency to human bone, heart, and other mesoderm cell types. Cell 166, 451–467.

26. Nakajima, T., Shibata, M., Nishio, M., Nagata, S., Alev, C., Sakurai, H., Toguchida, J., and Ikeya, M. (2018). Modeling human somite development and fibrodysplasia ossificans progressiva with induced pluripotent stem cells. Development 145, dev165431.

27. Nakajima, T., Nakahata, A., Yamada, N., Yoshizawa, K., Kato, T.M., Iwasaki, M., Zhao, C., Kuroki, H., and Ikeya, M. (2021). Grafting of iPS cell-derived tenocytes promotes motor function recovery after Achilles tendon rupture. Nature communications 12, 1–12.

28. Gentsch, G., Monteiro, R., and Smith, J. (2017). Cooperation between T-Box factors regulates the continuous segregation of germ layers during vertebrate embryogenesis. Current topics in developmental biology 122, 117–159.

29. Harvey, T., Flamenco, S., and Fan, C.-M. (2019). A Tppp3+ Pdgfra+ tendon stem cell population contributes to regeneration and reveals a shared role for PDGF signalling in regeneration and fibrosis. Nature cell biology 21, 1490–1503.

30. Shukunami, C., Takimoto, A., Nishizaki, Y., Yoshimoto, Y., Tanaka, S., Miura, S., Watanabe, H., Sakuma, T., Yamamoto, T., and Kondoh, G. (2018). Scleraxis is a transcriptional activator that regulates the expression of Tenomodulin, a marker of mature tenocytes and ligamentocytes. Scientific reports 8, 1–17.

31. Liu, H., Zhu, S., Zhang, C., Lu, P., Hu, J., Yin, Z., Ma, Y., Chen, X., and OuYang, H. (2014). Crucial transcription factors in tendon development and differentiation: their potential for tendon regeneration. Cell and tissue research 356, 287–298.

32. Takahashi, K., and Yamanaka, S. (2006). Induction of pluripotent stem cells from mouse embryonic and adult fibroblast cultures by defined factors. cell 126, 663–676.

33. Yamaguchi, T.P., Takada, S., Yoshikawa, Y., Wu, N., and McMahon, A.P. (1999). T (Brachyury) is a direct target of Wnt3a during paraxial mesoderm specification. Genes & development 13, 3185–3190.

34. Dunkman, A.A., Buckley, M.R., Mienaltowski, M.J., Adams, S.M., Thomas, S.J., Satchell, L., Kumar, A., Pathmanathan, L., Beason, D.P., and Iozzo, R.V. (2014). The tendon injury response is influenced by decorin and biglycan. Annals of biomedical engineering 42, 619–630.

35. Hofmann, M., Schuster-Gossler, K., Watabe-Rudolph, M., Aulehla, A., Herrmann, B.G., and Gossler, A. (2004). WNT signaling, in synergy with T/TBX6, controls Notch signaling by regulating Dll1 expression in the presomitic mesoderm of mouse embryos. Genes & development 18, 2712–2717.

36. Ishii, M., Arias, A.C., Liu, L., Chen, Y.-B., Bronner, M.E., and Maxson, R.E. (2012). A stable cranial neural crest cell line from mouse. Stem cells and development 21, 3069–3080.

37. Lin, H.-H., Bell, E., Uwanogho, D., Perfect, L.W., Noristani, H., Bates, T.J., Snetkov, V., Price, J., and Sun, Y.-M. (2010). Neuronatin promotes neural lineage in ESCs via Ca2+ signaling. Stem Cells 28, 1950–1960.

38. Zhao, X., and van Praag, H. (2020). Steps towards standardized quantification of adult neurogenesis. Nature Communications 11, 4275.

39. Yao, J.-j., Zhao, Q.-r., Lu, J.-m., and Mei, Y.-a. (2018). Functions and the related signaling pathways of the neurotrophic factor neuritin. Acta Pharmacologica Sinica 39, 1414–1420.

40. Berg, D.A., Su, Y., Jimenez-Cyrus, D., Patel, A., Huang, N., Morizet, D., Lee, S., Shah, R., Ringeling, F.R., and Jain, R. (2019). A common embryonic origin of stem cells drives developmental and adult neurogenesis. Cell 177, 654–668. e615.

41. Park, D.S., Seo, J.H., Hong, M., Bang, W., Han, J.K., and Choi, S.C. (2013). Role of Sp5 as an essential early regulator of neural crest specification in xenopus. Developmental Dynamics 242, 1382–1394.

42. Shi, J., Severson, C., Yang, J., Wedlich, D., and Klymkowsky, M.W. (2011). Snail2 controls mesodermal BMP/Wnt induction of neural crest. Development 138, 3135–3145.

43. Dehmelt, L., and Halpain, S. (2005). The MAP2/Tau family of microtubule-associated proteins. Genome biology 6, 1–10.

44. Simões-Costa, M., and Bronner, M.E. (2015). Establishing neural crest identity: a gene regulatory recipe. Development 142, 242–257.

45. Deak, K.L., Boyles, A.L., Etchevers, H.C., Melvin, E.C., Siegel, D.G., Graham, F.L., Slifer, S.H., Enterline, D.S., George, T.M., and Vekemans, M. (2005). SNPs in the neural cell adhesion molecule 1 gene (NCAM1) may be associated with human neural tube defects. Human genetics 117, 133–142.

46. Teppner, I., Becker, S., de Angelis, M.H., Gossler, A., and Beckers, J. (2007). Compartmentalised expression of Delta-like 1 in epithelial somites is required for the formation of intervertebral joints. BMC Developmental Biology 7, 1–12.

47. LeBlanc, L., Kim, M., Kambhampati, A., Son, A.J., Ramirez, N., and Kim, J. (2022). β-catenin links cell seeding density to global gene expression during mouse embryonic stem cell differentiation. Iscience 25, 103541.

48. Wu, J., Fan, Y., and Tzanakakis, E.S. (2015). Increased culture density is linked to decelerated proliferation, prolonged G1 phase, and enhanced propensity for differentiation of self-renewing human pluripotent stem cells. Stem cells and development 24, 892–903.

49. Kahane, N., and Kalcheim, C. (2020). Neural tube development depends on notochord-derived sonic hedgehog released into the sclerotome. Development 147, dev183996.

50. Piacentino, M.L., and Bronner, M.E. (2018). Intracellular attenuation of BMP signaling via CKIP-1/Smurf1 is essential during neural crest induction. PLoS Biology 16, e2004425.

51. Wymeersch, F.J., Wilson, V., and Tsakiridis, A. (2021). Understanding axial progenitor biology in vivo and in vitro. Development 148, dev180612.

52. Christ, B., Huang, R., and Scaal, M. (2004). Formation and differentiation of the avian sclerotome. Anatomy and embryology 208, 333–350.

53. Kult, S., Olender, T., Osterwalder, M., Markman, S., Leshkowitz, D., Krief, S., Blecher-Gonen, R., Ben-Moshe, S., Farack, L., and Keren-Shaul, H. (2021). Bi-fated tendon-to-bone attachment cells are regulated by shared enhancers and KLF transcription factors. Elife 10, e55361.

54. Blitz, E., Sharir, A., Akiyama, H., and Zelzer, E. (2013). Tendon-bone attachment unit is formed modularly by a distinct pool of Scx-and Sox9-positive progenitors. Development 140, 2680–2690.

55. Sugimoto, Y., Takimoto, A., Akiyama, H., Kist, R., Scherer, G., Nakamura, T., Hiraki, Y., and Shukunami, C. (2013). Scx+/Sox9+ progenitors contribute to the establishment of the junction between cartilage and tendon/ligament. Development 140, 2280–2288.

56. Miyabara, S., Yuda, Y., Kasashima, Y., Kuwano, A., and Arai, K. (2014). Regulation of Tenomodulin Expression Via Wnt/beta-catenin Signaling in Equine Bone Marrow-derived Mesenchymal Stem Cells. J Equine Sci 25, 7–13. 10.1294/jes.25.7.

57. Kishimoto, Y., Ohkawara, B., Sakai, T., Ito, M., Masuda, A., Ishiguro, N., Shukunami, C., Docheva, D., and Ohno, K. (2017). Wnt/beta-catenin signaling suppresses expressions of Scx, Mkx, and Tnmd in tendon-derived cells. PLoS One 12, e0182051. 10.1371/journal.pone.0182051.

58. Lara, H., Li, Z., Abels, E., Aeffner, F., Bui, M.M., ElGabry, E.A., Kozlowski, C., Montalto, M.C., Parwani, A.V., and Zarella, M.D. (2021). Quantitative image analysis for tissue biomarker use: a white paper from the digital pathology association. Applied Immunohistochemistry & Molecular Morphology 29, 479.

59. Schmittgen, T.D., and Livak, K.J. (2008). Analyzing real-time PCR data by the comparative C(T) method. Nat Protoc 3, 1101–1108. 10.1038/nprot.2008.73.

60. Jiang, W., Glaeser, J.D., Salehi, K., Kaneda, G., Mathkar, P., Wagner, A., Ho, R., and Sheyn, D. (2022). Single-cell atlas unveils cellular heterogeneity and novel markers in human neonatal and adult intervertebral discs. iScience 25, 104504. 10.1016/j.isci.2022.104504.

