## Supplement for "Single Cell Transcriptomics-Informed Induced Pluripotent Stem Cells Differentiation to Tenogenic Lineage"

**Suppl. Table 1:** Normalized cell counts (expressed as %) per cluster following WNTi treatment, shown for the two cell lines, 007i and 83i.

| **Cluster** | **Annotation** | **Markers** | **WNTi-007i**  **(% Normalized counts)** | **WNTi-83**  **(% Normalized counts)** |
| --- | --- | --- | --- | --- |
| **0** | SYN | MKX+TNMD+DCN+BGN | 66.85562 | 78.49919 |
| **1** | Mesoderm (Mes) | MIXL+TBXT+MSGN1+DLL1+DLL3+ | 3.40124 | 1.678803 |
| **2** | iPSC-1 | OCT4+,NANOG+LIN28A+SOX2+ | 3.418955 | 0.667476 |
| **3** | iPSC-2 | OCT4+,NANOG+LIN28A+SOX2+ | 2.710363 | 0.606796 |
| **4** | iPSC-3 | OCT4+,NANOG+LIN28A+SOX2+ | 3.790965 | 0.52589 |
| **5** | NMP/NC | TBXT/TWIST1/SP5/SNAI2 | 0.318866 | 0.101133 |
| **6** | iPSC-4 | OCT4+LIN28A+SOX2+ | 2.409212 | 2.04288 |
| **7** | iPSC-5 | OCT4+NANOG+LIN28A+SOX2+ | 3.525244 | 8.555825 |
| **8** | Neural crest | NTRK2+SOX4+SOX11+ | 4.69442 | 7.038835 |
| **9** | Fibrocartilage (FC) | COL2A1+SOX9+FN1+BGN+COL1A1 | 8.733392 | 0.262945 |
| **10** | Neural Lineage (NL) | SOX2+DCX+MAP2+UNCX+SOX4+ | 0.053144 | 0 |
| **11** | iPSC-6 | OCT4+NANOG+LIN28A+SOX2+ | 0.088574 | 0.020227 |

High Density Seeding

Low Density Seeding

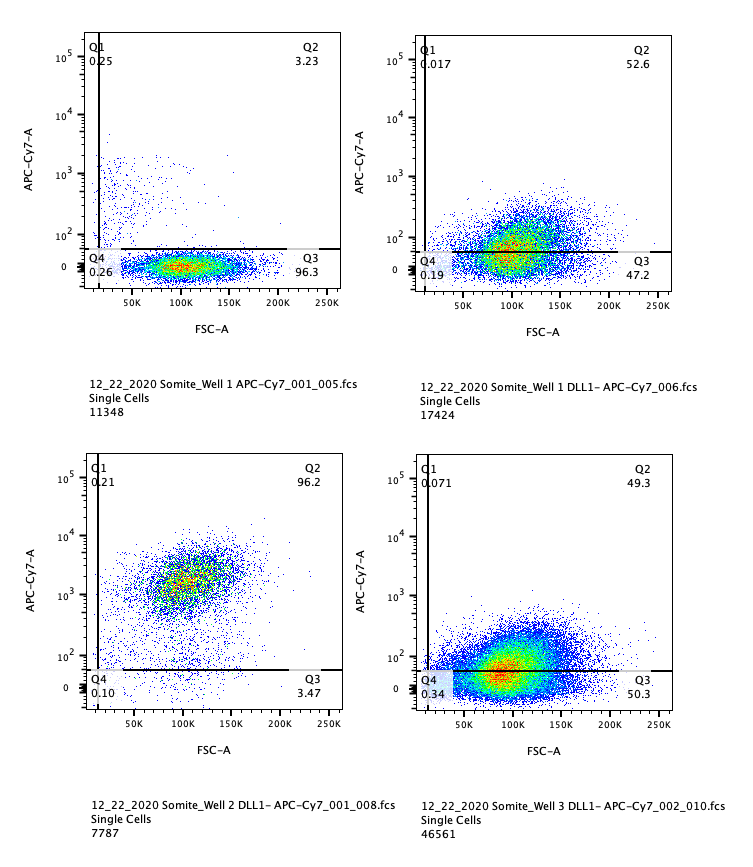

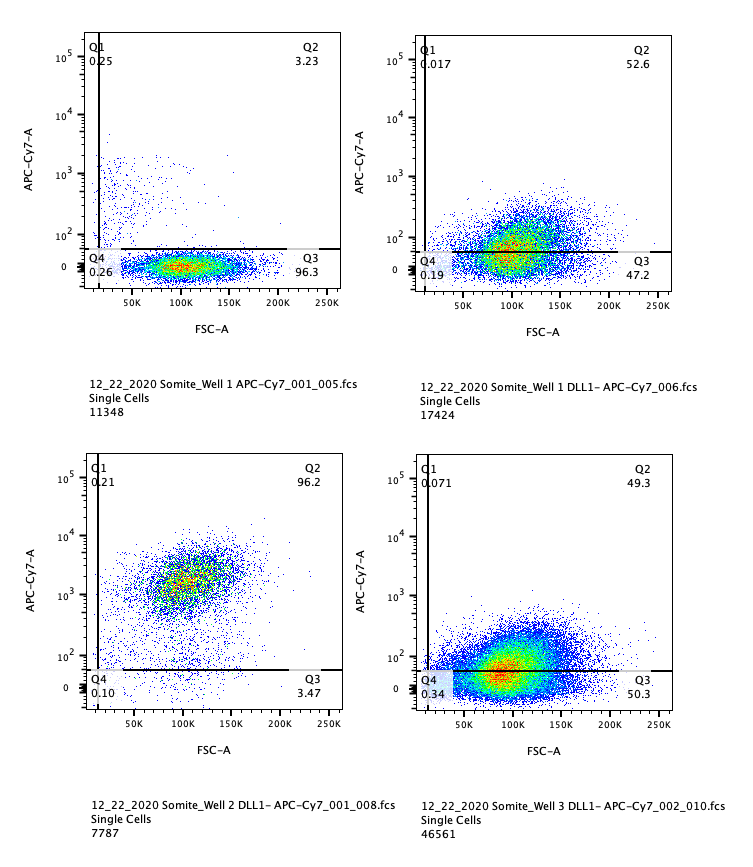

FSC-H/A

FSC-H/A

DLL-1

DLL1

**Suppl. Fig 1.** **Flow cytometry of DLL-1 at the PSM stage for low and high seeding densities.** **Left panel**: low iPSC seeding density resulted in high percentage of DLL1+ cells. **Right panel**: high iPSC seeding density resulted in a comparably reduced DLL1+ cell population.

Distribution of cell subpopulations

|  | iPSC | PSM | SM | SCL | SYN |
| --- | --- | --- | --- | --- | --- |
| iPSC-1 | 0.355878 | 0.171696 | 0.17137 | 0.171657 | 0.0529673 |
| iPSC-2 | 0.228137 | 0.148081 | 0.232055 | 0.215546 | 0.0840266 |
| iPSC-3 | 0.34531 | 0.147917 | 0.120548 | 0.180066 | 0.0493622 |
| SYN | 0.000528402 | 0.0990489 | 0.0813699 | 0.0842904 | 0.476428 |
| NMP/NC | 0.00409511 | 0.1653 | 0.0487671 | 0.134537 | 0.13117 |
| Mes | 0.000528402 | 0.115284 | 0.14863 | 0.09516 | 0.00277316 |
| iPSC-4 | 0.0521797 | 0.0995408 | 0.00780822 | 0.0223544 | 0.00804215 |
| iPSC-5 | 0.00779392 | 0.0455887 | 0.0660274 | 0.0451189 | 0.00970605 |
| NC | 0.000264201 | 0.00131191 | 0.117945 | 0.0168171 | 0.00277316 |
| NMP | 0.00462351 | 0.00541161 | 0.00534247 | 0.0246103 | 0.0715474 |
| Neural | 0.000660502 | 0.000819941 | 0.000136986 | 0.00984413 | 0.111204 |

**Suppl. Fig 2.** **Distribution of cell subpopulations per sample.** Cell clusters from iPSC to SYN were annotated to 6 distinct cell populations: iPSC (OCT4^+^SOX2^+^NANOG^+^), Syndetome (SYN, MKX^+^TNMD^+^COL1A1^+^), Neuromesodermal Progenitors/Neural Crest (NMP/NC, PAX3^+^NRP2^+^COLEC12^+^), Mesoderm (Mes, DLL1^+^DLL3^+^PARAXIS^+^), Neuromesodermal Progenitors – Cranial (NMP-C, DLL1^+^DLL3^+^NOTCH1^+^CRAB1^+^), and Neural Lineage cells (NL, NRN1^+^DCX^+^NNAT^+^).

A

007i SYN^WNTi^	83i SYN^WNTi^

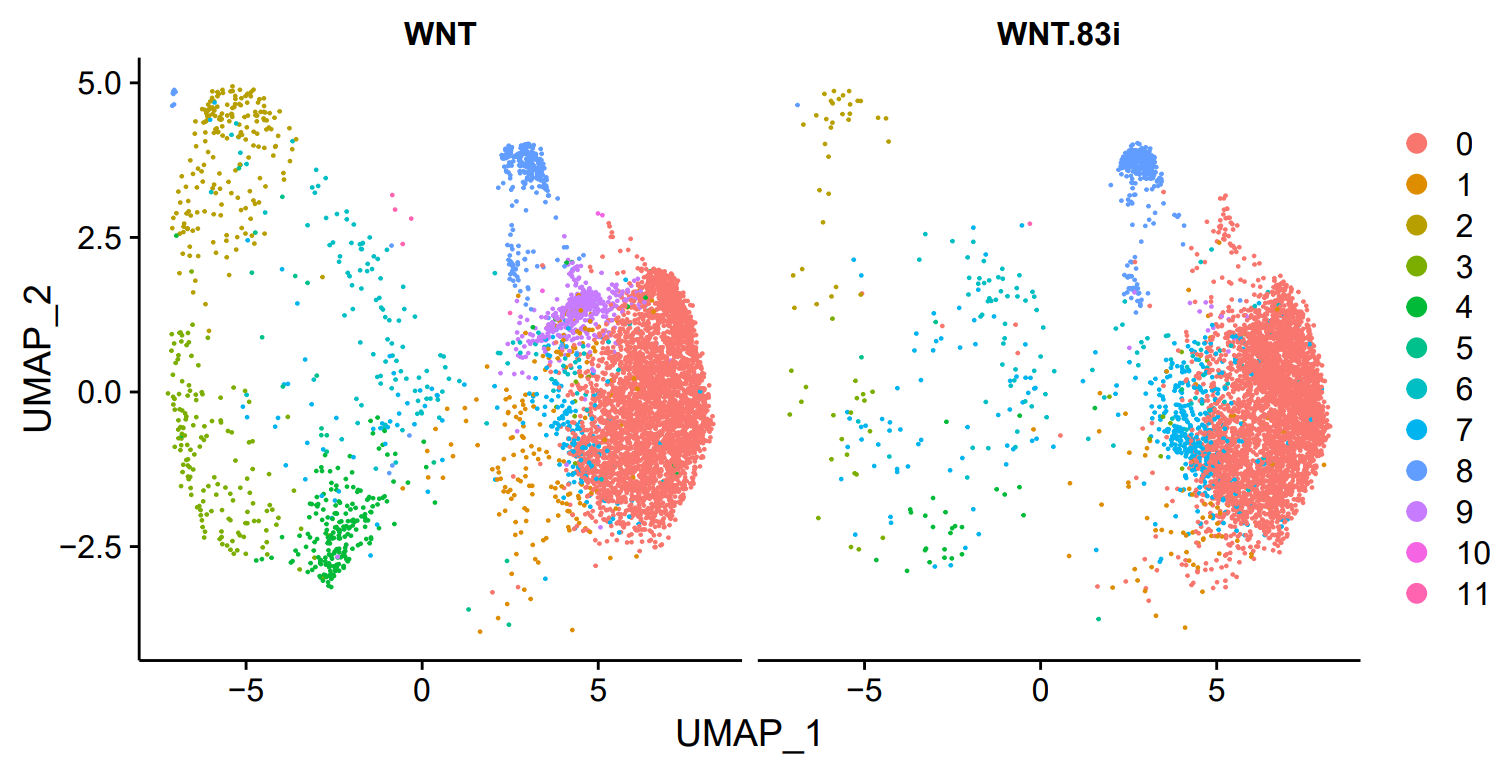

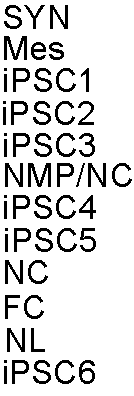

B

|  | 0 | 1 | 2 | 3 | 4 | 5 | 6 | 7 | 8 | 9 | 10 | 11 |
| --- | --- | --- | --- | --- | --- | --- | --- | --- | --- | --- | --- | --- |
| iPSCs | 0.198151 | 1.03038 | 33.144 | 28.4941 | 10.9247 | 2.39102 | 11.6909 | 9.39234 | 0.409511 | 0 | 0.0660502 | 2.25892 |
| PSM | 1.98426 | 27.4352 | 15.1689 | 15.0869 | 9.6589 | 15.8413 | 10.6592 | 3.62414 | 0.491965 | 0 | 0.0491965 | 0 |
| SM | 0.150685 | 25.0959 | 11.9041 | 14.8219 | 19.6438 | 18.0411 | 9.10959 | 0.643836 | 0.452055 | 0 | 0.0136986 | 0.123288 |
| SCL | 1.39487 | 24.7179 | 16.9026 | 15.6308 | 15.5077 | 12.1846 | 9.25128 | 0.697436 | 2.17436 | 0.0205128 | 1.10769 | 0.410256 |
| SYN | 27.5374 | 34.0544 | 4.85302 | 5.15807 | 5.87909 | 1.19246 | 2.6345 | 0.277316 | 6.04548 | 0.110926 | 12.2573 | 0 |
| WNTi-007i | 66.8556 | 3.40124 | 3.41895 | 2.71036 | 3.79097 | 0.318866 | 2.40921 | 3.52524 | 4.69442 | 8.73339 | 0.0531444 | 0.088574 |
| WNTi-83i | 78.4992 | 1.6788 | 0.667476 | 0.606796 | 0.52589 | 0.101133 | 2.04288 | 8.55583 | 7.03883 | 0.262945 | 0 | 0.0202265 |

**Suppl. Fig. 3.** ***A).*** ScRNA-seq comparison of two different iPSC lines for SYN^WNTi^. Cell population annotation is shown in Fig.5 and in Suppl. Table 1. ***B).*** Distribution of cell subpopulations per sample, where the second cell line sample at the syndetome stage has also been included (designated as WNTi-83i). Clusters numbers and annotations are shown in Suppl. Table 1.

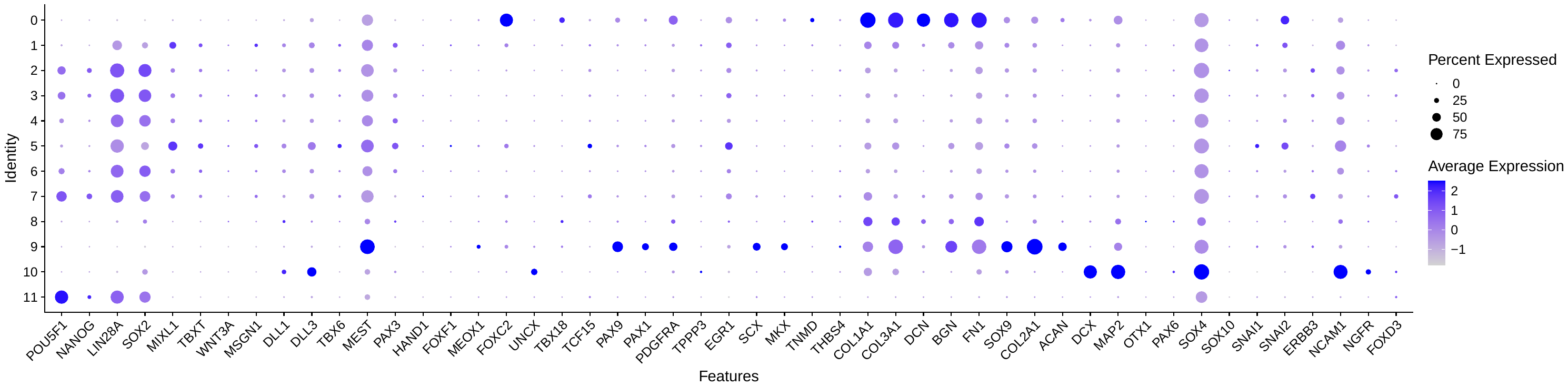

**Suppl.Fig. 4**. Dot plot of stage-specific markers of SYN^WNTi^ for each cluster. Cell population annotation is presented in Fig.5.

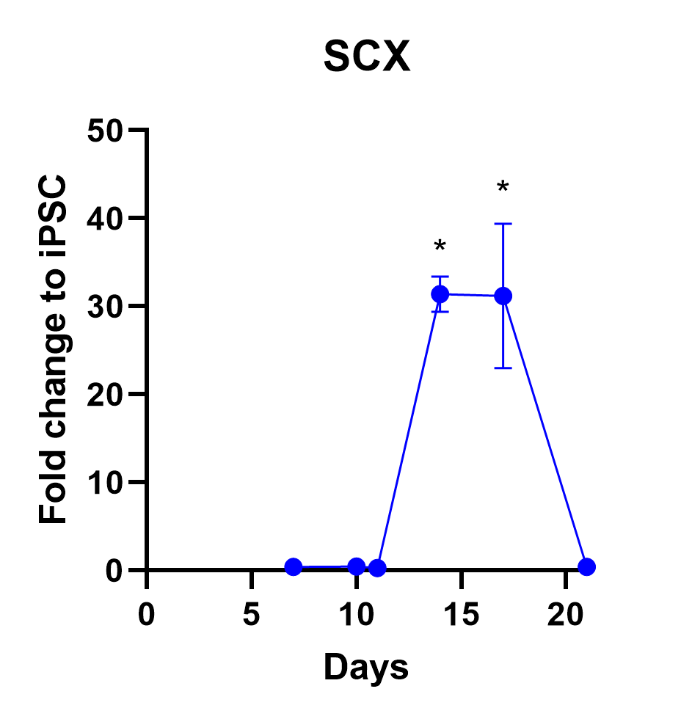

**Suppl. Fig. 5.** Change in SCX expression throughout SYN induction with WNTi. All timepoints are normalized to day 0 (iPSC). *p<0.0001.

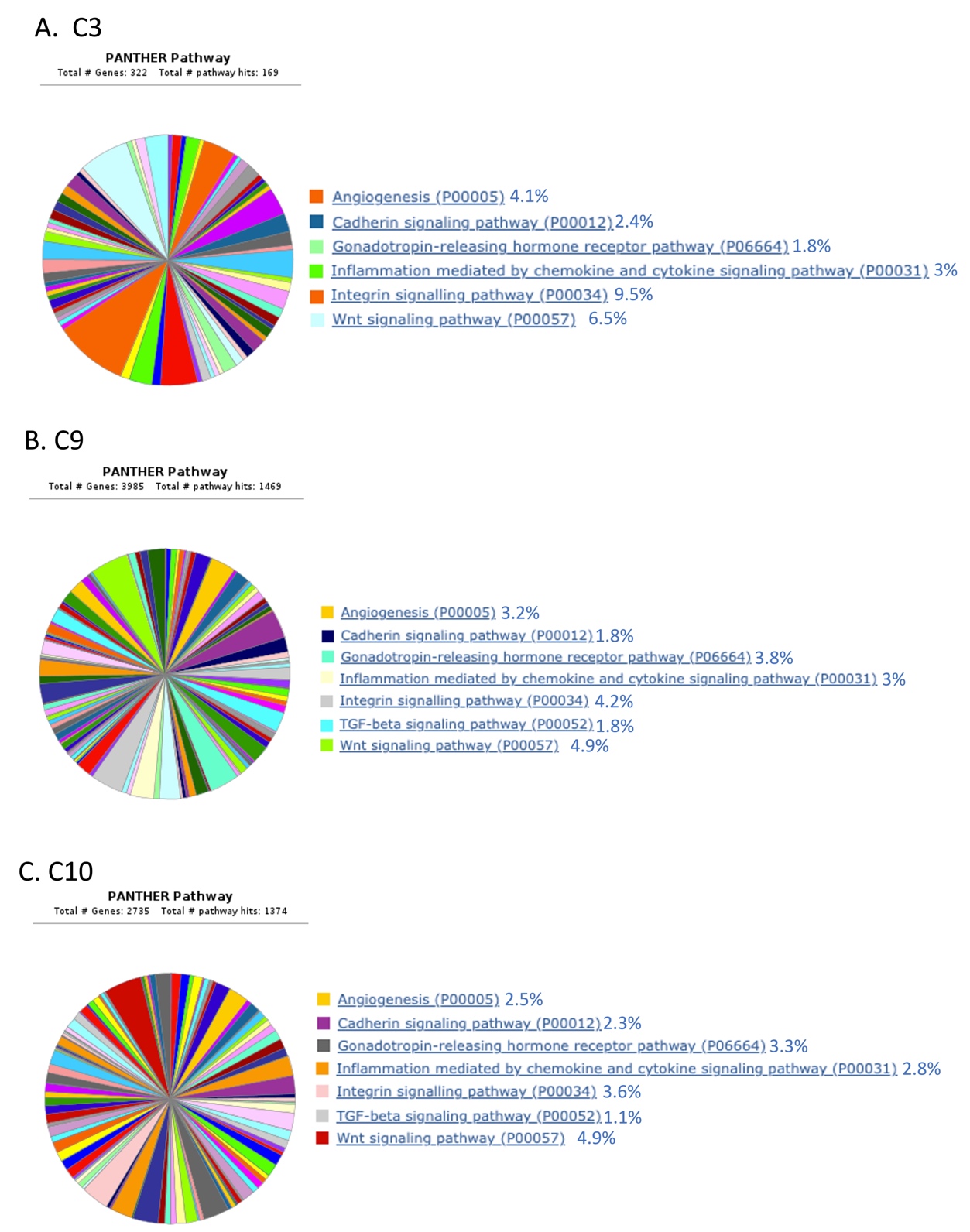

**Suppl. Fig.6.** A non-biased GO analysis was performed. Multiple pathways were detected in the three pathways of interest, that is C3 (Fig. 3A; SYN), C9 (Fig. 3A; NMP/NC) and C10 (Fig. 3A; NL).

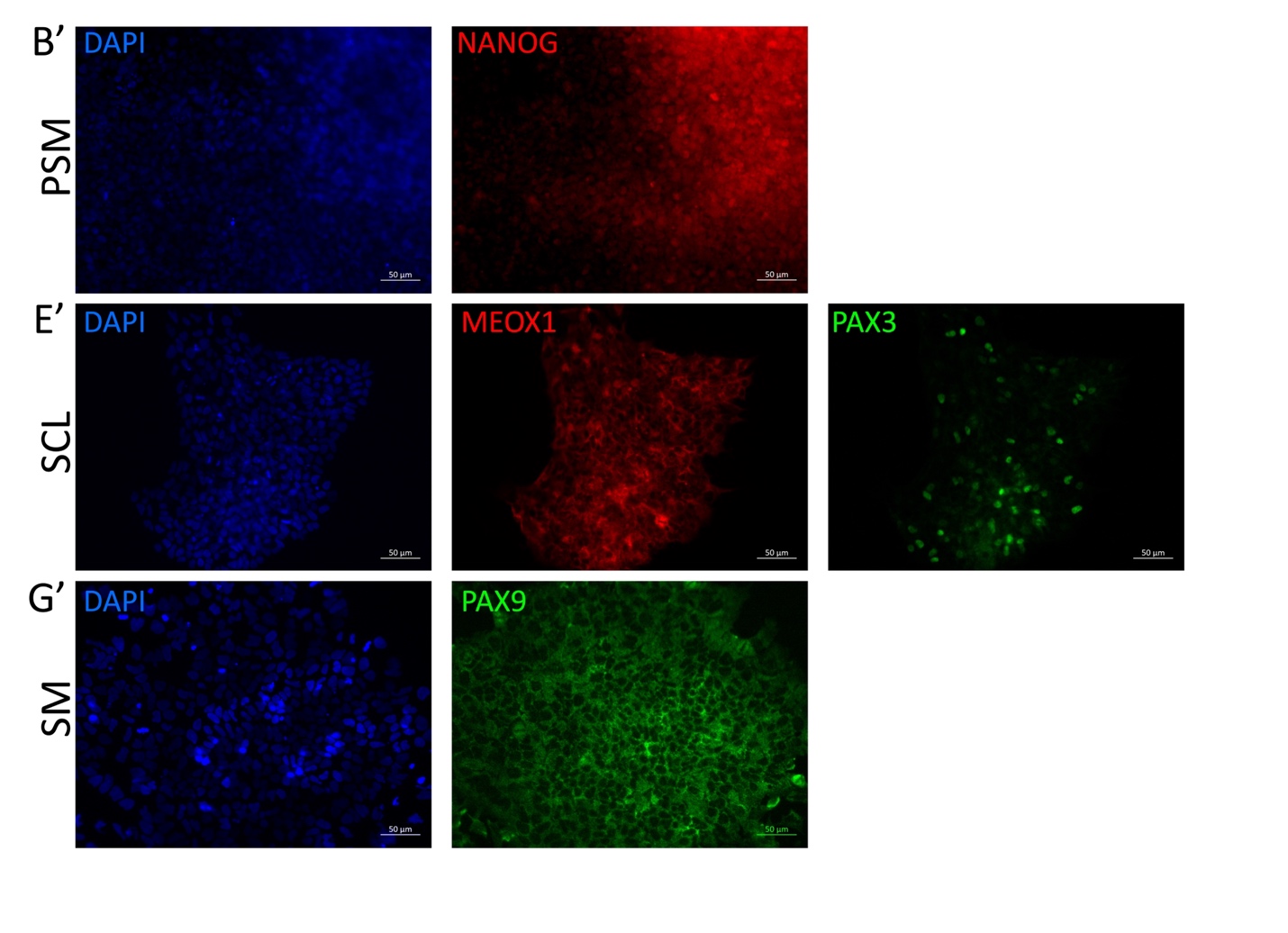

**Suppl. Fig. 7.** Immunofluorescence staining of selected markers and DAPI for nuclear staining from Fig.2 expanded to show all separate channels. (B’) NANOG at PSM stage; (E’), MEOX1 and PAX3 at SCL stage (G’) PAX9 at SM stage.

SCL

SYN

SYN-WNTi

**Suppl. Fig. 8.** Separate and combined feature plots of SCX and TNMD expression at SCL through SYN stage with and without WNT inhibitor (WNTi).

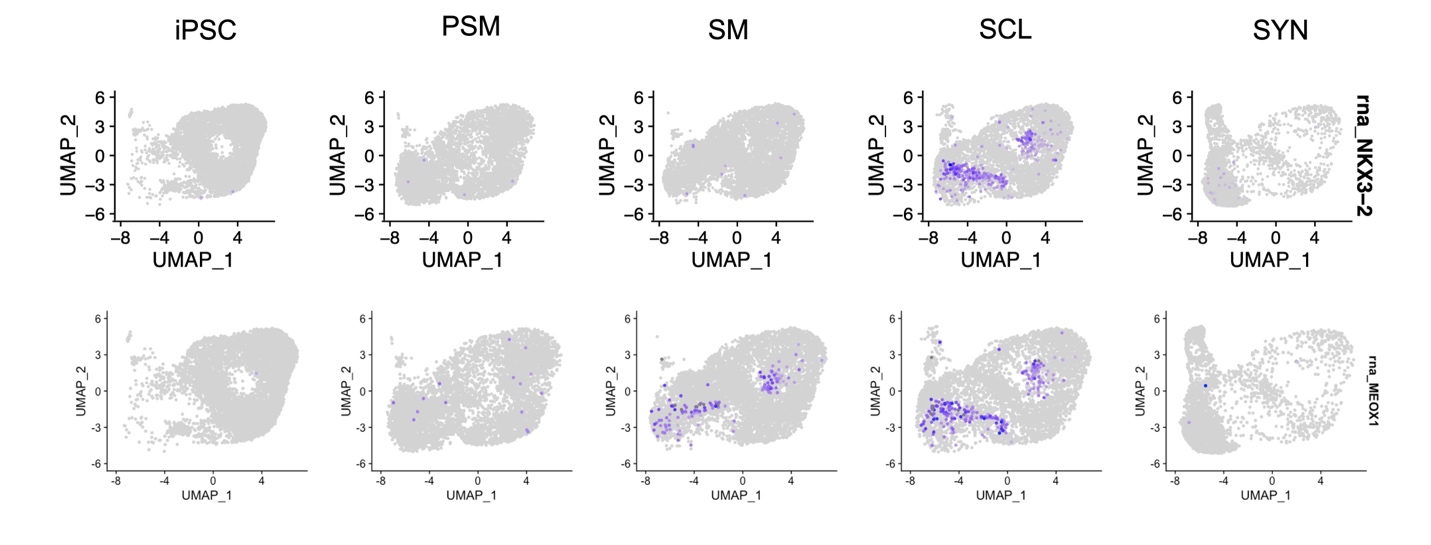

**Suppl. Fig. 9.** Feature plots of NKX3.2 and MEOX1 displayed for all differentiation stages.
